# Glia multitask to compensate for neighboring glial cell dysfunction

**DOI:** 10.1101/2024.09.06.611719

**Authors:** Allison N. Beachum, Gabriela Salazar, Amelia Nachbar, Kevin Krause, Hannah Klose, Kate Meyer, Ariana Maserejian, Grace Ross, Hannah Boyd, Thaddeus Weigel, Lydia Ambaye, Hayes Miller, Jaeda Coutinho-Budd

## Abstract

As glia mature, they undergo glial tiling to abut one another without invading each other’s boundaries. Upon the loss of the secreted neurotrophin Spätzle3 (Spz3), *Drosophila* cortex glia transform morphologically and lose their intricate interactions with neurons and surrounding glial subtypes. Here, we reveal that all neighboring glial cell types (astrocytes, ensheathing glia, and subperineurial glia) react by extending processes into the previous cortex glial territory to compensate for lost cortex glial function and reduce the buildup of neuronal debris. However, the loss of Spz3 alone is not sufficient for glia to cross their natural borders, as blocking CNS growth via nutrient-restriction blocks the aberrant infiltration induced by the loss of Spz3. Surprisingly, even when these neighboring glia divert their cellular resources beyond their typical borders to take on new compensatory roles, they are able to multitask to continue to preserve their own normal functions to maintain CNS homeostasis.

**Graphical Abstract:** 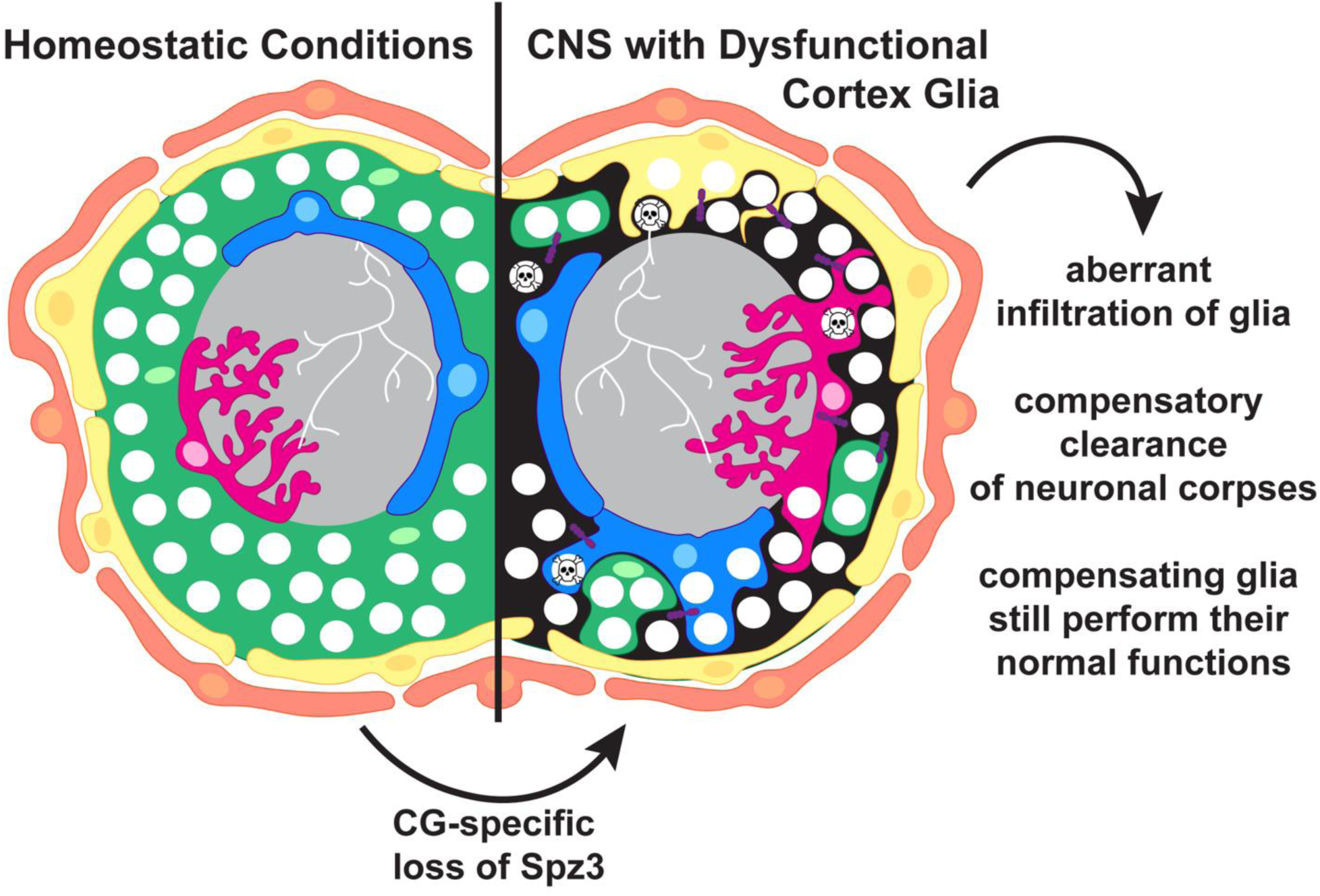

## Introduction

Glia play active roles in setting up and maintaining the nervous system, including ongoing sculpting of the nervous system throughout development, homeostasis, and disease.^1–5^ Beyond their key interactions with neuronal counterparts, glial cells can influence each other both directly and indirectly. For example, glia can be directly linked by gap junctions that allow for fast communication between glia cells of the same subtype, such as astrocytes in a wide-spanning network.^6^ Alternatively, glial cells can interact across subtypes, such as microglia secreting factors to activate neurotoxic astrocytes or astrocytes directly coupled to oligodendrocytes by gap junctions.^7,8^ Within the same glial subtype, glial cells undergo a phenomenon known as tiling, where each glial cell grows to maximize contact with the enveloped neuronal counterpart while minimizing contact with their glial neighbors. Glial tiling exists throughout the animal species from worms^9^ and flies,^10,11^ to zebrafish,^12^ mice,^13–16^ and humans.^17^ The size of the domains and overlap varies between subtype, species, age, and nervous system health,^17–20^ yet the conserved recurrence of glial tiling from species to species argues for its importance. However, limited knowledge exists about these glial-glial interactions: How are they built and maintained? What roles do these interactions have in supporting central nervous system (CNS) homeostasis?

In vertebrates, glial tiling is typically studied within a single glial subtype, such as the stereotyped spacing of cell bodies and minimally-overlapping domains of astrocytes,^12,13,21^ microglia,^22,23^ or oligodendrocyte precursor cells;^24^ however, intermingling among glial cells of different subtypes makes it difficult to fully understanding how glial cells of different subtypes respond to altered tiling or general dysfunction of another glial type. Like the vertebrate system, the fly CNS is composed of multiple neuronal and glial subtypes that work in concert to maintain nervous system function. In particular, these cells share many morphological, cellular, genetic, and functional features with their mammalian counterparts.^25,26^ Additionally, the relative simplicity of the invertebrate nervous system coupled with powerful genetic tools allows for unparalleled depth of investigation^27–29^ that can be used to easily genetically label and manipulate distinct cell types in multiple ways simultaneously.

The spatial organization of the *Drosophila* CNS proves to be particularly useful in the context of glial tiling. Not only do *Drosophila* glial cells tile within subtypes, such as astrocytes to astrocytes, but they form tiled boundaries with neighboring glial subtypes as well (Figure 1A). Specifically, the *Drosophila* CNS is composed of multiple morphologically distinct classes of glial cells that are compartmentalized across two primary regions: the cortex that consists of neuronal cell bodies and their proximal axons and the neuropil containing all the CNS neurites and synapses.

**Figure 1.**
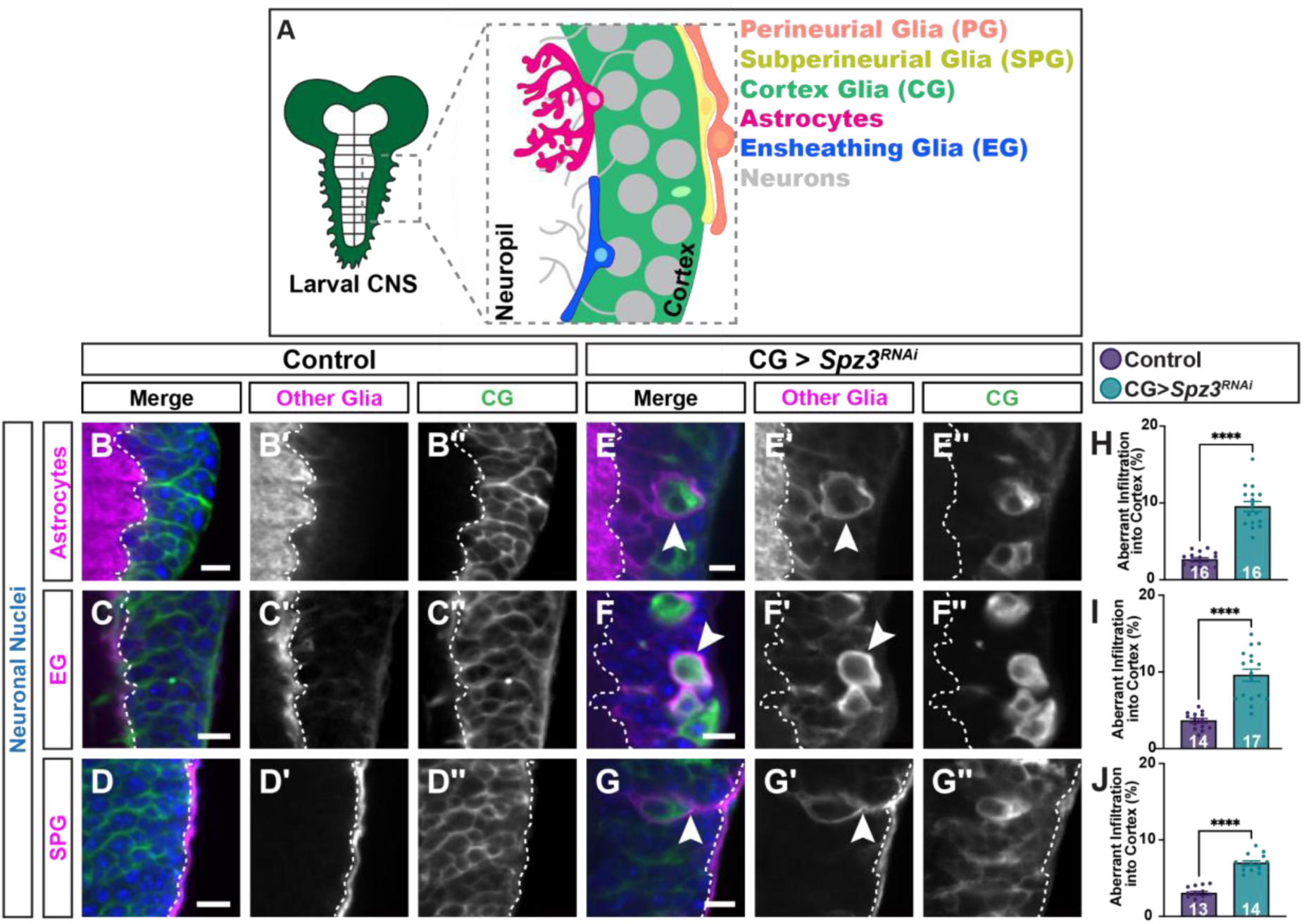
Neighboring glia infiltrate the cortex when CG morphology is altered. (**A**) The *Drosophila* central nervous system (CNS) is composed of a neuronal cell cortex (dark green) and the synaptic neuropil (white). The cortex houses neuronal somas (gray), cortex glia (light green; CG), and the the cell bodies of neighboring glial subtypes such as astrocytes (pink) and ensheathing glia (blue; EG). The CNS synaptic contacts are found within the neuropil (white) alongside the astrocyte processes, which fill up the neuropil. EG (blue) line the cortex-neuropil boundary. Perineurial glia (orange) and the subperineurial glia (yellow; SPG) surround the nervous system and make up the BBB. (**B-G**) Control brains (B-D) with CG (green) and neuronal nuclei (anti-Elav, blue) show normal CG in the VNC encasing each neuronal cell body with neighboring astrocytes (B), EG (C), and SPG (D) depicted in magenta demonstrating stereotyped localization under homeostatic conditions, minimally infiltrating the cortex. (E-G) CG expressing RNAi against Spz3 (Spz3^RNAi^) results in globular morphology, allowing all neighboring glia to infiltrate into the cortex (astrocytes in E, EG in F, and SPG in G). White dashed line denotes the boundary between the cortex and neuropil regions of the VNC. White arrowheads denote areas of aberrant infiltration. Scale bars = 10 µm. (**H-J**) Quantification of aberrant infiltration of astrocytes (H), EG (I), and SPG (J) into the cortex. N = number of animals, denoted in each corresponding bar. (****) *P* < 0.0001. Unpaired t-test.

Importantly, glia in the *Drosophila* CNS are arranged along these compartmental boundaries, with individual subtypes covering specific regions with minimal overlap with their neighbors. Starting from the CNS surface, the entire nervous system is covered by a thin layer of perineurial glia (PG) cells that serve as one of the first boundaries between the surrounding hemolymph and the neurons. Located just underneath the PG cells are the aptly named subperineurial glia (SPG), the main components that form and maintain the blood-brain barrier (BBB) in *Drosophila.*^28,30–33^ SPG are flat cells that form strong intercellular boundaries with elaborate septate junctions to prevent paracellular diffusion.^30,34,35^ Underneath the SPGs, cortex glia (CG) are housed within the cortex, weaving thin cellular processes amidst the neuronal cell bodies in a lace-like manner to support the somas.^25,31^ A single CG cell interacts with an average of 50 neuronal cell bodies in the larva^36^ to provide key metabolic support^37^ and clear neuronal debris as the CNS develops and ages.^4,36^ CG span from the SPGs on the surface to both types of neuropil associated glia: ensheathing glia (EG) and astrocytes. EG form a sheath around the neuropil, helping to separate this region from the cell body cortex. Astrocyte cell bodies are located along the cortex-neuropil border, and their extensive network of arborized processes invade the neuropil to help modulate synaptic transmission and maintenance.^21,25^ Each of these subtypes tile with cells of their own kind to fill their designated regions, while other glial subtypes are excluded from these domains.^10,33,34,36^

Tiled adjacent glial cells have been shown to react to their compromised neighbors of the same subtype. For instance, upon astrocyte ablation or the weakening of astrocytes by genetic mutations, any remaining astrocytes will extend their domain coverage within the neuropil to minimize the detrimental effect on synaptic regulation.^21,38^ Similarly, when a single CG cell is ablated, neighboring CG move in to ensure that neuronal cell bodies are properly wrapped.^36^ We recently showed that this space-filling response is not confined to cells of the same subtype. During the onset of CG disruption, astrocytic processes begin to invade the cortex.^11,36^ Whether this reaction signifies a specific relationship between CG and astrocytes or if this reactivity applies more broadly to all neighboring glial cells has remained unresolved. Tangentially, EG ablation has been shown to result in compensatory proliferation of astrocytes to offset the loss of their neuropil glial counterpart,^10^ but whether this proliferation is a damage response or a general mechanism to account for glial compensation is unclear. Furthermore, these cross-glial interactions are not merely a phenomenon seen in invertebrate biology. In mammalian systems, microglia primarily localize to and remove apoptotic neuronal cell bodies, while astrocytes localize to more distal neuronal process to clean up dying neurites and synaptic debris^39^ – paralleling the relationship between CG and astrocytes in the *Drosophila* CNS. Upon the loss of soma-associated microglia, astrocytes extend processes towards caspase+ cell bodies and upregulate lysosomal activity, albeit at a less efficient rate than microglia.^39^ However, it is unclear how universally-applicable this phenomenon is. Moreover, if glial cells react to impaired neighboring glial cells by diverting their resources to carrying out a compensatory role for another glial cell such as in response to dysfunctional glia that give rise to glioma,^40^ how well can they maintain their own functions?

In the current study, we investigated multiple glial-glial interactions and, following the morphological disruption of CG via CG-specific loss of Spz3, and found aberrant outgrowth into the cortex during development from all glial subtypes with direct CG interactions. Specifically, in addition to astrocytes, EG and SPG readily extend cellular processes into the cortex. Surprisingly, even with the constitutive knockdown of Spz3, restricting animal growth prevented both the CG disruption and infiltration from other glial cell types, suggesting that this neighboring glial infiltration is not simply due to reduced Spz3 signaling but also the loss of CG function as the CNS grows. While loss of Spz3 impairs CG proliferation in addition to morphological maintenance, our data reveal that proliferation is not necessary for maintaining full CG coverage throughout the cortex. Furthermore, driving proliferation in the Spz3-depleted cells is not sufficient to rescue morphology and full coverage of CG processes throughout the cortex. We show that when CG- somal interactions are disrupted, astrocytes, EG, and SPG all contribute to the clearance of caspase+ neuronal cell bodies to compensate for lost CG function. However, compensatory infiltration changes throughout lifespan, as ensheathing glia become the predominant infiltrating cell type for cortex glial dysfunction in the adult CNS. Perhaps most remarkably, even when astrocytes, SPG, and EG divert their cellular resources outside of their normal boundaries to take on these new compensatory roles, they are still capable of performing their own homeostatic functions of synaptic maintenance (astrocytes), post-injury clearance (EG), and BBB maintenance (SPG). This work illustrates that surrounding, presumably healthy glial cells can react to nearby glial dysfunction in other subtypes by extending into their region and multitasking to preserve proper nervous system function and maintain CNS homeostasis.

## Results

### All neighboring glial subtypes infiltrate into the cortex when CG morphology is altered

Under normal conditions, CG are the primary glial cells positioned in the cortex to interact with neuronal cell bodies. Our previous studies demonstrated that when CG are morphologically transformed from lacelike to globular cells by the CG-specific knockdown of Spz3 (Spz3^RNAi^; Figure 1E-G), astrocyte processes aberrantly infiltrate into the cortex (Figure 1E’).^11,36^ Here, we have implemented enhanced analytical methods to more thoroughly assess the extent of this infiltration in three dimensions with precise quantification. A small number of astrocyte processes were occasionally observed extending short distances into the cortex of control animals, occupying less than 3% of the total cortex volume (Figure 1B’,H; 2.62% ± 0.24% of cortex area). In contrast, there was a significant increase in the amount of aberrant infiltration of astrocytic processes into the cortex in the presence of globular CG and unwrapped neurons (Figure 1E’,H; 9.53% ± 0.67%; 3.63-fold increase). As astrocytes are not the only neighboring glial subtype to tile with CG, we wondered if this relationship was specific to astrocyte-CG interactions, or whether other glial cells that share a boundary with CG might exhibit similar reactions. In control brains, EG line the boundary between the cortex and neuropil regions (Figure 1C’) and SPG surround the outside perimeter of the CNS (Figure 1D’). Like astrocytes, EG (Figure 1C,I) and SPG (Figure 1D,J) have been recently reported to occasionally extend small processes short distances into the cortex under normal conditions.^41^ Here, we quantified the extent of these EG and SPG projections in control animals, which invade only a small percentage of the cortex volume (3.64% ± 0.29% and 3.03% ± 0.25%, respectively); however, in the presence of transformed CG, there was a significant increase in the amount of aberrant infiltration of both EG (Figure 1F,I; 9.57% ± 0.79%; 2.67-fold increase) and SPG (Figure 1G,J; 6.98% ± 0.30%; 2.31-fold increase) into the cortex. These observations show that the presence of morphologically and functionally intact CG restricts the growth of astrocyte, EG, and SPG processes to their respective domains. Moreover, these data demonstrate that a singular genetically-compromised glial subtype can affect the growth and reactivity of multiple neighboring glia.

### Growth restriction blocks CG alteration and neighboring glia infiltration

In Spz3^RNAi^ animals, CG are established properly during embryonic stages and remain morphologically intact through 2nd instar larval (L2) stages, but are dramatically altered to form the globular phenotype during the mid to late larval period (L2-L3).^36^ During the L2 to L3 transition, quiescent neuroblasts are reactivated by nutrient-dependent signals to begin to produce more neurons that will eventually compose the mature adult CNS.^42^ Throughout this same period, CG undergo endomitosis to rapidly increase their number of nuclei, becoming multinucleated cells, and extend processes to keep up with this sudden CNS growth and enwrap the newly born neurons.^36,43^ We hypothesized that the onset of this rapid growth period could trigger the morphological CG transformation induced by the loss of Spz3. To test this, we performed a growth restriction assay (Figure 2A) where 1st instar larvae (L1) were selected and transferred to one of two conditions: 1) normal molasses food for 5 days (until they reach L3) or 2) 20% sucrose agar for the same time period. The sucrose agar deprives the larvae of key nutrients required for animal growth, ensuring that the animals are age-matched with congruent RNAi expression for the duration of the experiment; however, animals raised on 20% sucrose agar remain the size of L1 animals, preventing the exponential growth that occurs between L2-L3. When raised on molasses food and allowed to grow to L3, control animals (Figure 2B) exhibit full CG coverage that enwrap nearly all neuronal cell bodies, but Spz3-depleted CG adopt the globular morphology (Figure 2C) as shown previously. However, in animals raised on 20% sucrose agar that remain at L1 for the duration, both the control (Figure 2D) and Spz3^RNAi^ (Figure 2E) animals preserve normal cortex glial morphology and coverage (Figure 2L). These data indicate that the loss of CG morphology at L2 is triggered by the onset of exponential growth at this time.

**Figure 2.**
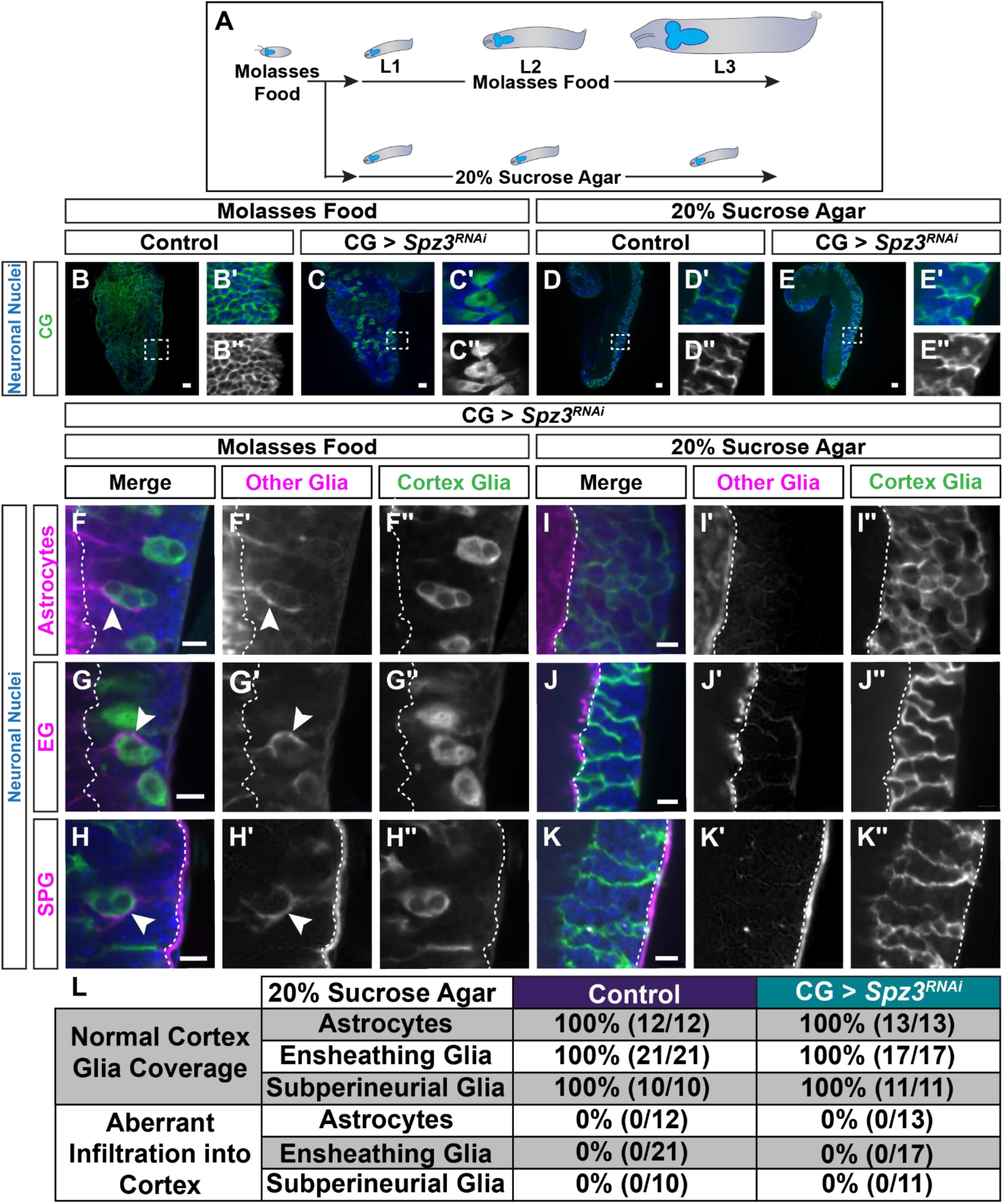
Nutrient-deprived growth restriction blocks CG transformation and neighboring glia infiltration. (**A**) Animals raised on molasses food progress through the three larval stages normally during which time the body and CNS grow exponentially (top). To restrict growth of the larvae, L1s are transferred to a 20% sucrose agar, aged out 5 more days, during which time the animals’ bodies or CNSs do not grow in size (bottom). (**B-C**) Control animals raised on molasses food progress through larval stages, and CG (green) wrap each neuronal cell body (blue) for a full coverage of the cortex (B). Spz3^RNAi^ in CG results in progressive morphological disruption of CG by the L3 stage (C). (**D-E**) In animals raised on 20% sucrose agar, CG morphology remains intact (D) and Spz3^RNAi^ no longer results in disrupted CG morphology (E). Scale bars = 20 µm (B-E). (**F-K**) Age-matched Spz3^RNAi^ animals raised on molasses food reach L3, and demonstrate disrupted CG and aberrant infiltration of the neighboring glia into the cortex (F-H). Spz3^RNAi^ animals raised on 20% sucrose agar with (I) astrocytes (magenta), (J) EG (magenta), and (K) SPG (magenta) labeled demonstrates that without CNS growth, there is normal CG morphology (F”-K”) and no aberrant infiltration of neighboring glia (F’-K’). White dashed line denotes the boundary between the cortex and neuropil regions of the VNC. White arrowheads denote areas of aberrant infiltration. Scale bars = 10 µm (F-K). (**L**) Quantification of animals with normal cortex glial morphology and coverage, and whether aberrant infiltration from neighboring glia was observed in control and Spz3^RNAi^ knockdown animals.

To determine if it was the knockdown of Spz3 or the alteration of CG morphology that was required for neighboring glia infiltration, we observed the morphology and territories of astrocytes, EG, and SPG in Spz3^RNAi^ animals raised on molasses food or 20% sucrose agar. As shown in Figure 1, when Spz3^RNAi^ animals are raised on molasses food, CG are morphologically disrupted and astrocytes (Figure 2F’), EG (Figure 2G’), and SPG (Figure 2H’) infiltrate into the cortex. In Spz3^RNAi^ animals raised on 20% sucrose agar, where RNAi is still expressed but the CG are not morphologically altered, neighboring glia no longer infiltrate into the cortex (Figure 2I-L). Together these data suggest that the exponential growth that occurs between L2-L3 is required for both the disruption of CG morphology and subsequent infiltration of neighboring glia into the cortex in Spz3^RNAi^ animals, rather than the loss of Spz3 alone.

### Proliferation is not necessary or sufficient to rescue disrupted CG morphology

When CG fail to maintain their lacelike morphology and adopt a globular morphology as result of Spz3^RNAi^ expression, they also fail to proliferate as they should between L2-L3,^36^ leading us to question whether the disruption of proliferation was linked to the morphological transformation and whether one might directly cause the other. To determine if blocking CG proliferation could recapitulate the Spz3-regulated phenotype, we expressed RNAi targeting *string (stg)* specifically in CG to inhibit proliferation by blocking the transition into M-phase,^44^ and quantified the number of CG nuclei. Compared to controls (Figure 3A,D; 231.35 ± 9.40), Stg knockdown effectively inhibits proliferation of CG in the thoracic segments of the VNC, as evidenced by the reduced number of CG nuclei (Figure 3B,D; Stg^RNAi-17760^: 81.53 ± 7.07 and Stg^RNAi-34831^: 44.65 ± 1.13). CG in the abdominal segments do not normally proliferate between L2-L3,^36^ and therefore Stg knockdown does not reduce the numbers of CG nuclei in the abdominal segments (Figure 3D; Control: 51.05 ± 1.33; Stg^RNAi-17760^: 49.58 ± 2.25 and Stg^RNAi-34831^: 50.62 ± 1.33). Strikingly, CG expressing Stg^RNAi^ still extend processes to fully wrap neurons throughout the cortex despite the dramatically reduced nuclear number (Figure 3B’). To confirm this finding, we also tested RNAi targeting *Cyclin A (CycA)*–an additional protein needed for cell cycle progression–specifically in CG and quantified the number of CG nuclei (Figure S1A-B). Similarly, knocking down CycA decreases the number of CG nuclei while maintaining wild-type CG morphology and territory throughout the cortex (Figure S1C; CycA^RNAi-35694^: 99 ± 2.86 and CycA^RNAi-103595^: 113 ± 5.86). These data demonstrate that CG proliferation is not necessary to maintain proper CG growth, morphology, territory, or interactions with neuronal cell bodies.

**Figure 3.**
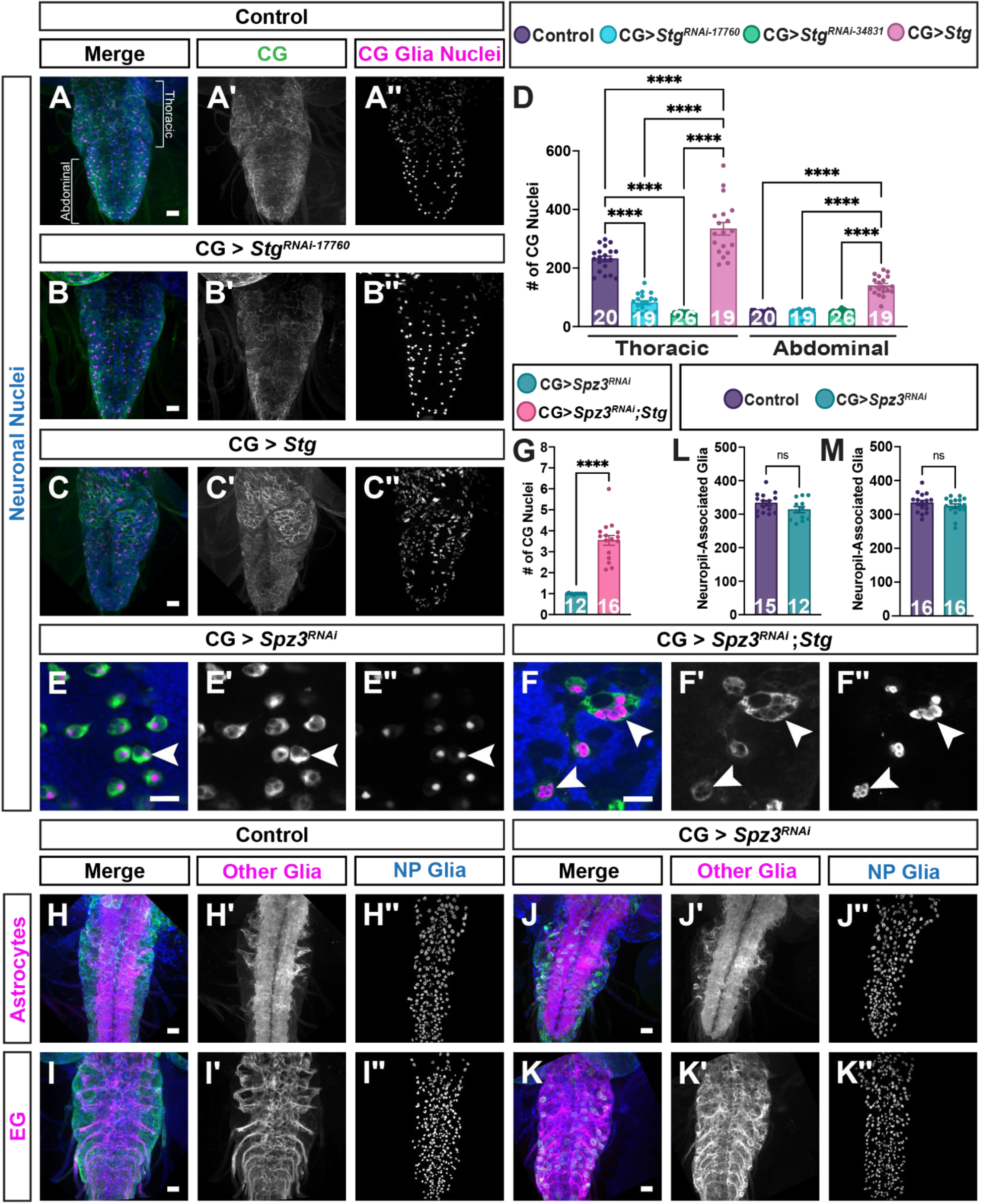
Proliferation is not necessary or sufficient to maintain CG morphology, and neuropil-associated glial number doesn’t change in response to disrupted CG. (**A**) Control VNCs with CG processes (green), CG nuclei (magenta) and neuronal nuclei (anti-Elav, blue) depict normal CG throughout the VNC. (**B**) CG expressing RNAi against Stg inhibits CG proliferation resulting in morphologically normal CG, but a decreased number of CG nuclei. (**C**) Overexpressing Stg in CG drives proliferation and increases the number of CG nuclei. (**D**) Quantification of CG nuclei. (****) *P* < 0.0001, One-Way ANOVA with multiple comparisons. N = number of animals, denoted inside each corresponding bar. (**E-F**) CG loss of Spz3 disrupts CG morphology and number of nuclei (E), but driving proliferation by Stg overexpression in Spz3 knockdown CG is not sufficient to rescue CG morphology (F). White arrowheads denote globular CG with one nuclei (E) or multiple nuclei (F). (**G**) Quantification of the number of CG nuclei per globular morphology. (****) *P* < 0.0001, Unpaired t-test. (**H**) Control and (**J**) CG-specific Spz3 knockdown (green) with genetically-labeled astrocytes (magenta) with, and all glial nuclei (anti-repo, blue). (**I**) Control and (**K**) Spz3 knockdown in CG (green) in the EG- labeled condition (EG, magenta), and glial nuclei (anti-repo, blue). Grayscale images depict astrocytes (H’) or EG (K’) and the neuropil-associated Repo+ glial nuclei (H’’-K’’). Scale bars = 20 µm (E-F) or 50 µm (A-C, H-K). (**L-M**) Quantification of neuropil-associated glial (NP) nuclei. N = number of animals, denoted inside corresponding bars. (n.s.) Not significant. Unpaired t- test.

Conversely, to determine whether extensive proliferation would disrupt CG growth and morphology, we overexpressed Stg in CG. While this expression was sufficient to increase the number of CG nuclei in the VNC thoracic segments (Figure 3C-D; 334.11 ± 21.32) as well as in the typically non-proliferative CG of the abdominal segments (Figure 3C-D; 139.89 ± 7.46), CG still maintained full cortex coverage and intact morphology (Figure 3C’). To determine whether driving proliferation in the Spz3-depleted CG could rescue morphology, we simultaneously knocked down Spz3 while overexpressing Stg in CG. Stg expression still led to increased proliferation in these cells, as we observed a greater number of nuclei per globular CG (Figure 3E-G; Control: 1.00 ± 0.01; Spz3^RNAi^+Stg: 3.51 ± 0.27), but ultimately did not rescue the overall morphology of the CG nor their ability to wrap neurons (Figure 3F’). These data further support our finding that CG proliferation, morphology, and infiltration throughout the cortex are not directly coupled.

### Astrocyte and EG do not proliferate or migrate to infiltrate the cortex in response to CG disruption

Although CG proliferation is not necessary to maintain proper CG morphology and somal interactions, it remained unclear whether the aberrant infiltration from neighboring glia was due merely to process extension from existing cells, or whether these glia proliferate or migrate in order to compensate for the loss of CG coverage. Glial proliferation has been demonstrated in response to CNS damage,^45^ and astrocytes have been shown to proliferate after EG ablation,^10^ so we sought to examine whether a similar phenomenon occurred in response to disrupted CG. As SPG are multinucleated glia, we chose to focus this assessment on the uninucleate neuropil-associated glia. Astrocytes and EG in the larval VNC are large cells with nuclei located at very stereotyped positions around the larval neuropil. To determine whether the number of neuropil-associated glia is altered in Spz3^RNAi^ animals, we visualized both subsets of glial nuclei with Repo (Figure 3), a pan-glial nuclear marker, or Prospero (Figure S2) which specifies astrogenic programming protein that can differentiate astrocyte and EG nuclei at the neuropil.^46^

We assessed control and Spz3^RNAi^ animals at L3, after the onset of CNS growth and disruption of CG morphology, while labeling each neuropil glia subset; however, to better label each nucleus and to ensure additional specification we visualized glial nuclei with anti-Repo and anti-Prospero in both the astrocyte (Figure 3H,J, S2E,G) and EG (Figure 3I,K, S2F,H) driver backgrounds. Compared to the number of Repo+ neuropil-associated glial nuclei in control animals (astrocytes: 333.2 ± 7.56, EG: 334.25 ± 7.09), Spz3^RNAi^ animals did not demonstrate a statistically significant difference in the number or location of neuropil-associated glial nuclei (Figure 3L-M; astrocytes: 313.67 ± 9.17, EG: 324.44 ± 6.83). Similarly, quantifying the number of Prospero+ nuclei in both the control astrocyte (Figure S2A,E; 157.88 ± 4.29) and EG (Figure S2B,F; 149.50 ± 2.91) driver backgrounds demonstrates there was no difference in the number or location of Prospero+ neuropil-associated glia when there is aberrant infiltration in the cortex (Figure S2C,D; Astrocytes: 160.29 ± 4.16, EG: 148.06 ± 2.53). These data suggest that the infiltrating neighboring glia do not increase in cellular number nor do they migrate into the cortex in order to take over cortex glial space. Instead, these data support the fact that the infiltrating glia send out additional processes to extensively infiltrate into the cortex.

Repo also labels a population of proliferating cells that line the neuropil of the thoracic segments of the L3 VNC. These proliferating neuropil glia give rise to EG and motor neurons of the adult CNS.^47^ These cells are labeled by both our astrocyte and EG drivers, but they are not established astrocytes or EG at this stage of development so we therefore quantified this population separately. Interestingly, there is no difference in the number of proliferating neuropil glia in the astrocyte background (Figure S2G,I,K; control: 147.26 ± 4.17, Spz3^RNAi^: 139.25 ± 11.91), but there is a decrease in the EG background (Figure S2H,J,L; control: 178.80 ± 9.58, Spz3^RNAi^: 115.15 ± 7.23). Additionally, we often found that these proliferating cells appeared to have migrated away from the neuropil in Spz3^RNAi^ animals (Figure S2M,N), which could subsequently impact the future lineage of these cells, including mature EG in the adult.

### Neighboring glial processes clear neuronal debris to compensate for lost CG function

The morphological transformation of CG due to Spz3 depletion results in increased caspase+ neuronal nuclei (Figure 4G-I).^36^ Whether this primarily increase results from the loss of nutrient support from CG or the inability for CG to clear neuronal corpses is unclear, as it is difficult to parse these apart when CG are no longer present to do either function. However, one of the major documented roles of CG is to clear neuronal cell corpses throughout development;^4,48–52^ therefore, the lack of CG interactions at neuronal cell bodies results in the accumulation of excessive neuronal debris. In morphologically intact CG, blocking their ability to engulf via the removal of the major glial engulfment receptor, Draper (Drpr, the homolog of mammalian MEGF10),^53^ results in a far greater number of lingering caspase+ neuronal nuclei.^4^ It is therefore possible that the neighboring glial cells infiltrate the cortex to clear neuronal debris in a compensatory fashion. Such compensation has been suggested between mammalian astrocytes and microglia after microglial ablation.^39^ To determine if all neighboring glia compensate for dysfunctional CG, we simultaneously blocked engulfment in each of the infiltrating glial cell types in the presence of CG-specific Spz3^RNAi^, and compared death caspase protein 1 (DCP-1) positive nuclei across four conditions for each glial subset. We first quantified DCP-1+ nuclei in each genetic driver control animals when astrocytes (Figure 4A’, 346.88 ± 14.35), EG (Figure 4B’, 197.71 ± 11.36), or SPG (Figure 4C’, 318.63 ± 8.07) are simply labeled with one fluorophore and CG are labeled with another. While CG remain intact, there is no statistically significant difference in the number of DCP1+ puncta present in the cortex when *drpr* is knocked down (Drpr^RNAi^) only in astrocytes (Figure 4D,M, 358.06 ± 13.66), EG (Figure 4E,O, 286.1% ± 14.60), or SPG (Figure 4F,Q, 379.55 ± 17.32) as these cells are not located in the cortex to perform this role under normal conditions. Alternatively, when Spz3^RNAi^ is expressed in CG without Drpr^RNAi^ in neighboring glia, as expected there is a significant increase in the number of DCP1+ puncta in multiple driver backgrounds: astrocytes (Figure 4G,M, 448.94 ± 27.07, 1.29-fold increase) and EG (Figure 4H,O, 452.24 ± 43.09, 2.29-fold increase), though the increase does not reach significance in the SPG background (Figure 4I,Q, 372.47 ± 25.61, 1.17-fold increase). Lastly, when Spz3^RNAi^ is expressed in CG and Drpr^RNAi^ is expressed in one neighboring glial subset, we observed a dramatic increase in the number of DCP1+ puncta in each: astrocytes (Figure 4J,M, 576.8 ± 40.90, 1.66-fold increase), EG (Figure 4K,O, 617.71 ± 54.75, 3.12-fold increase), and SPG (Figure 4L,Q, 489.00 ± 30.99, 1.53-fold increase). It is worth noting that reducing Drpr in the neighboring glia did not inhibit processes from infiltrating into the cortex (Figure 4N,P,R), so Drpr signaling does not appear to be necessary for calling the processes into the cortex. Together these results suggest that when CG morphology is disrupted and there are inadequate glial-somal interactions, neighboring glia infiltrate the cortex at least in part to clear additional neuronal corpses in compensation for the lack of CG function.

**Figure 4.**
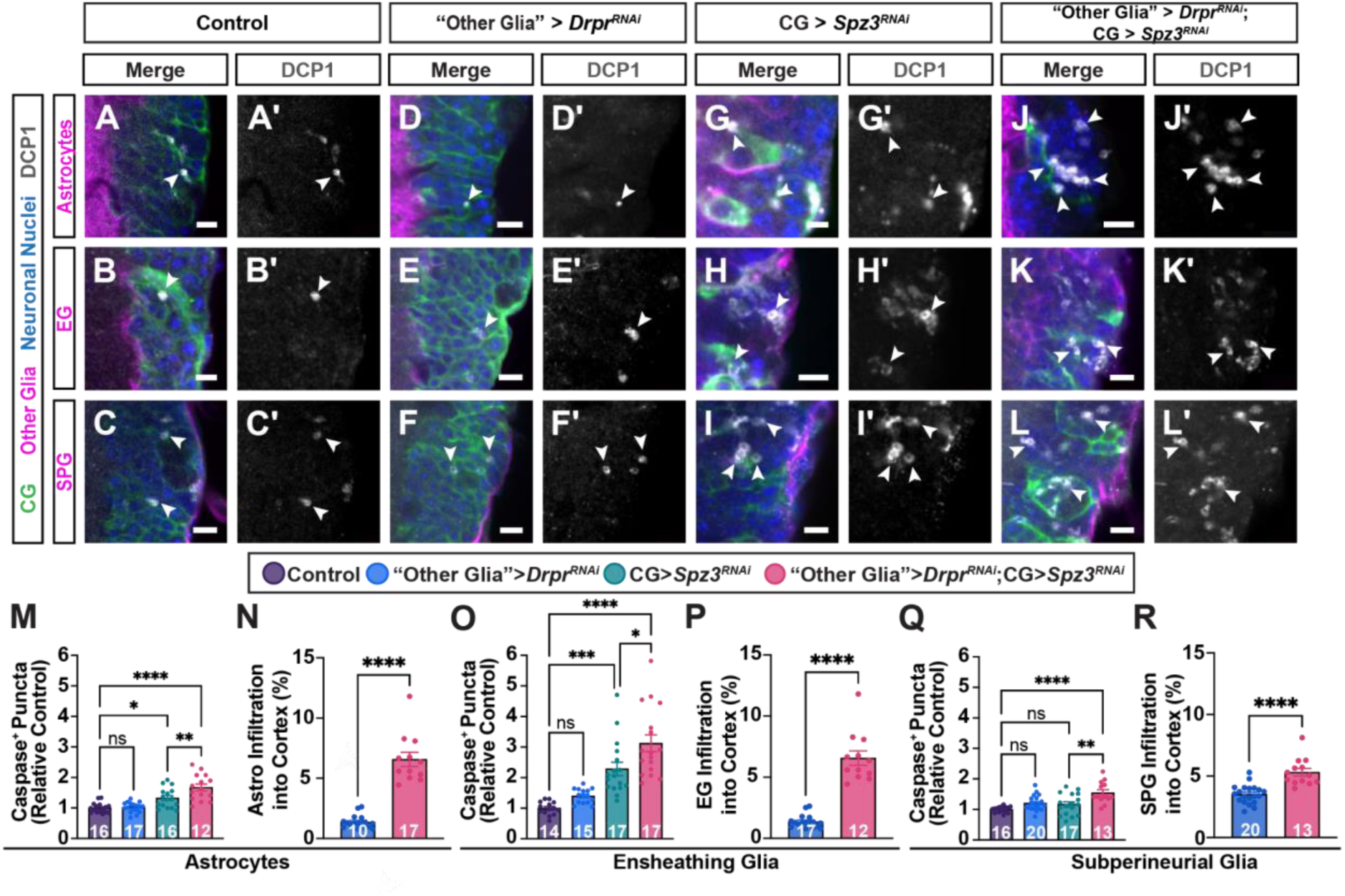
Aberrant glial processes compensate for CG function to clear out neuronal corpses. (**A-C**) Control VNC with astrocytes (A, magenta), EG (B, magenta), or SPG (C, magenta), CG (green), neuronal nuclei (anti-Elav, blue), and caspase+ nuclei (anti-DCP1, gray) show minimal levels of dying neurons in the cortex. (**D-F**) Drpr knockdown in astrocytes (D, magenta), EG (E, magenta), or SPG (F, magenta) does not alter the number of caspase+ cells when CG are morphologically normal. (**G-I**) When Spz3 is knocked down in CG, DCP-1+ cells increase. (**J-L**) When Drpr is knocked down in astrocytes (J), EG (K), or SPG (L), and Spz3 is simultaneously knocked down in CG, the number of DCP1+ cells increases even further. Grayscale images depict the DCP1 channel alone (A’-L’). Scale bars = 10 µm. White arrowheads denote DCP-1+ puncta. (**M, O, Q**) Quantification of the number of DCP1+ puncta in the cortex of the VNC. (**N, P, R**) Quantification of the aberrant infiltration of each neighboring glial subtype into the cortex, depicted as ratio to the control for each respective driver background. N = number of animals, denoted inside corresponding bars. (n.s.) Not significant. (*) *P* < 0.5, (**) *P* < 0.01, (***) *P* < 0.001, (****) *P* < 0.0001. One-Way ANOVA with multiple comparisons (M, O, Q). Unpaired T-test (N, P, R).

### Prolonging the presence of dying neurons increases neighboring glial infiltration when CG morphology is altered

We have shown that the aberrantly invading glial processes can function to remove extra neuronal debris when cortex glia cannot; however, whether cues from these dying neurons aid in calling additional processes into the region or if the glial processes grow into the area and then subsequently perform roles once they are there was unclear. To determine if dying neuronal cues might signal to neighboring glial processes to call them into the cortex and aid in engulfment, we drove cell death in neurons with the pro-apoptoic gene *rpr*^54^ or blocked the final steps of neuronal cell death with *p35,* a baculovirus gene that inhibits downstream effector caspases such as DrICE and DCP-1.^55,56^ Overexpression of rpr under the control of a neuronal-specific driver was confirmed by qPCR (Figure S3A) and expression of p35 was confirmed via a p35-specific antibody (Figure S3B-C). When used in conjunction with Spz3^RNAi^ in CG, we observed no statistical difference in the percentage of glial infiltration between controls (Figure 5A,D 3.65 ± 0.44%) compared to neuronal rpr expression (Figure 5B,D, 2.34% ± 0.36%). Strikingly, in animals expressing Spz3^RNAi^ in CG and an overexpression of p35 in neurons, we saw a significant increase in the percentage of glial infiltration (Figure 5C,D, 6.48% ± 0.87%). Of note, signaling of the apoptotic pathway upstream of these effector caspases is still intact and now prolonged in these cells^57^ implying that one of these upstream signaling factors could result in a “find me” cue to call in neighboring glial processes in the absence of functional cortex glia. Together these data demonstrate that neighboring glia infiltrate the cortex in the presence of globular CG to clear additional neuronal debris in response to signals released from dying neurons.

**Figure 5.**
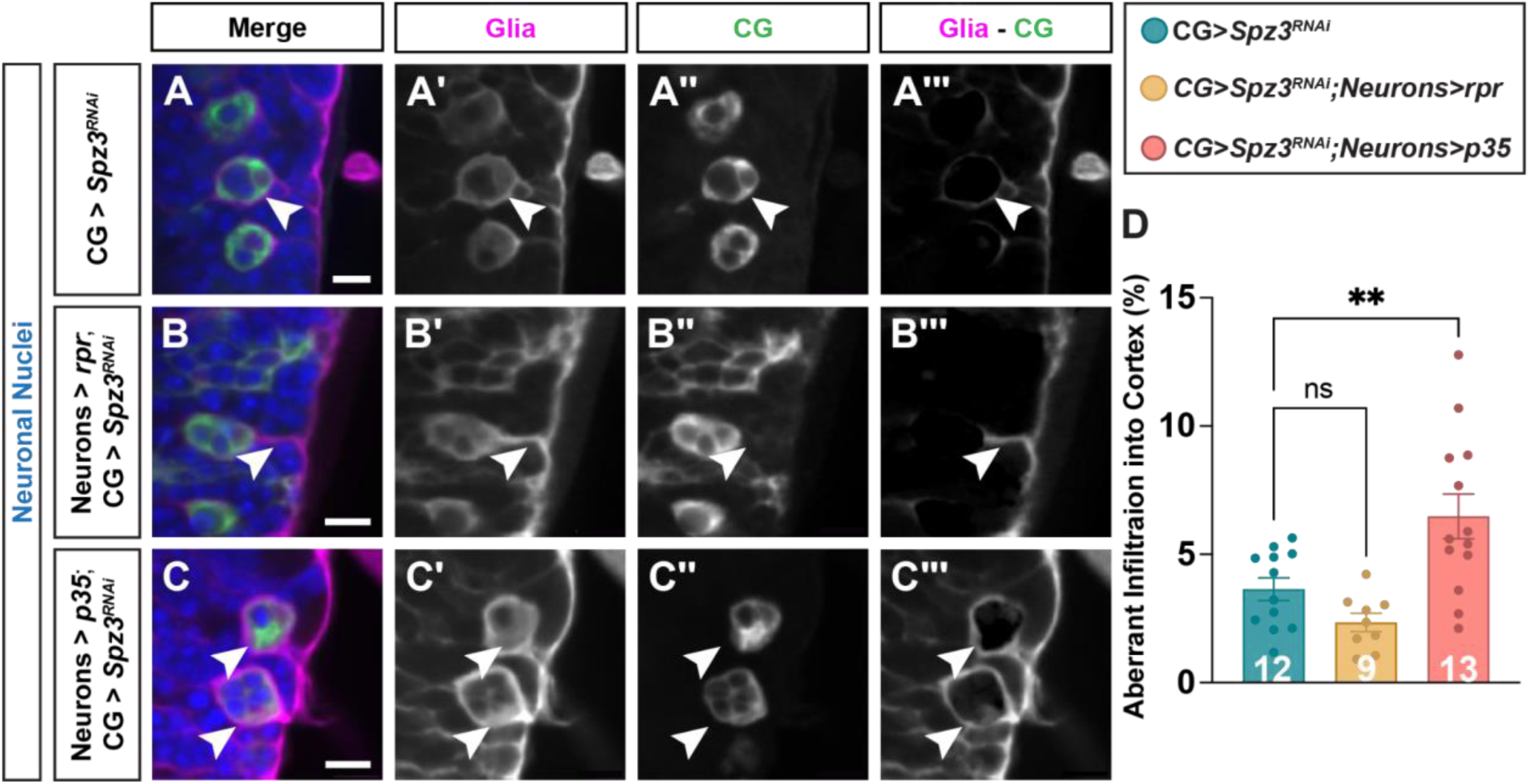
Blocking the execution of cell death increases aberrant glial infiltration when CG morphology is altered. (**A-C**) Spz3^RNAi^ in CG with CG (green), glia (3xP3-RFP labeling multiple glial subtypes, magenta), and neuronal nuclei (anti-Elav, blue) demonstrates aberrant infiltration of glia into the cortex (A). The addition of neuronal overexpression of rpr to increase neuronal cell death (B) results in no significant change in glial infiltration, while overexpression of p35 to inhibit neuronal cell death (C) results in a significant increase in neighboring glial infiltration into the cortex. Grayscale images of total glial channel (A’-C’), CG channel (A’’-C’’), or the aberrant infiltration of non-CG alone (A’’’-C’’’). Scale bars = 10 µm. White arrowheads denote areas of aberrant infiltration. (**D**) Quantification of the aberrant infiltration into the cortex. N = number of animals, denoted inside corresponding bars. (n.s.) Not significant. (*) *P* < 0.5, (**) *P* < 0.01, (***) *P* < 0.001, (****) *P* < 0.0001. One-Way ANOVA with multiple comparisons.

### In the absence of CG, EG are the primary infiltrating glial cell type in the adult VNC

While the CNS compartmentalization remains the same between larval and adult stages (Figures 1A, 6A), metamorphosis brings about a profound reorganization of the circuitry during this time (Figure 6A), not only for neurons but also for glia.^47,58–60^ However, many details regarding how glial cells of the CNS behave throughout metamorphosis remain unresolved. SPGs of the CNS persist from embryonic to adult stages and do not divide during metamorphosis;^25,34^ however, larval astrocytes and EG undergo programmed cell death during mid metamorphosis.^49,60^ The neuropil-associated glial cells that reside in the adult brain are derived from multipotent Type II neuroblast lineages during late larval and early pupal stages.^47,60,61^ Given the restructuring of the CNS throughout this later developmental period, we wanted to investigate how the continued loss of Spz3 in CG affected glial-glial interactions with the neighboring glial cells in adulthood.

**Figure 6.**
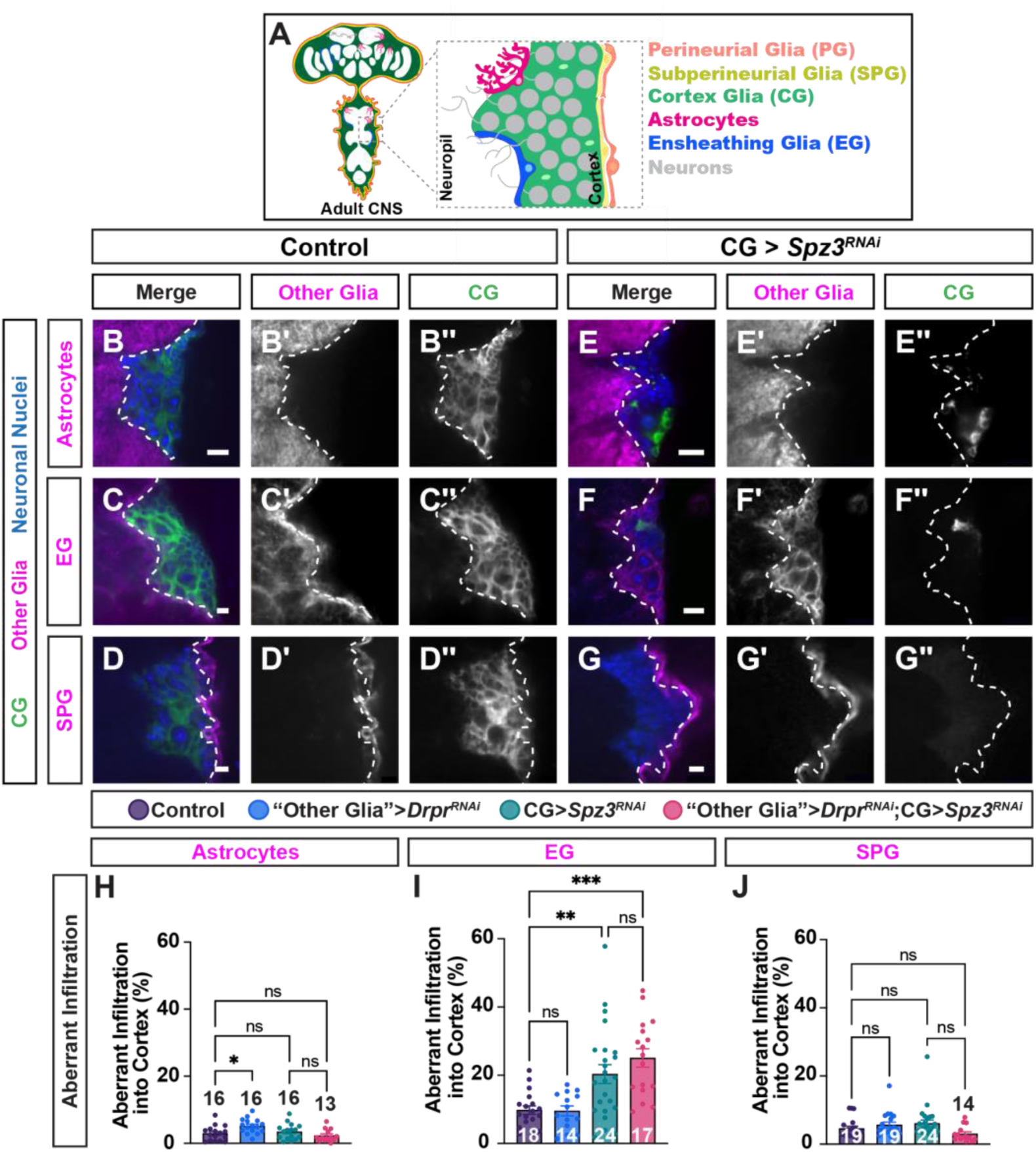
EG are the primary infiltrating glial cell type in the adult VNC. (**A**) Diagram of the Drosophila adult CNS morphology and inset of the VNC depicting the imaged region in B-G. (**B-D**) Control adults show normal CG (green) in the VNC encasing neuronal cell bodies (anti-elav, blue) with astrocytes (B), EG (C), and SPG (D) in magenta depicting stereotypical localization with minimal infiltration into the cortex. (**E-G**) CG expressing Spz3^RNAi^ are largely depleted from the adult VNC, allowing neighboring EG the ability to infiltrate into the cortex (F), while astrocytes (E) and SPG (G) do not. White dashed line denotes the boundary between the cortex and neuropil. Scale bar = 10 µm. (**H-J**) Quantification of aberrant infiltration of (H) astrocytes, (I) EG, and (J) SPG into the cortex. N = numbers of animals, denoted in each corresponding bar. (n.s.) Not significant. (*) *P* < 0.5, (**) *P* < 0.01, (***) *P* < 0.001. Two-Way ANOVA with multiple comparisons.

To determine if the predominant phenotypes observed in CG Spz3^RNAi^ L3 animals persisted into the adult, we investigated (1) the amount of aberrant infiltration performed by neighboring glia (Figure 6), (2) the number of neuropil-associated glial nuclei (Figure S4), and (3) the whether infiltrating neighboring glia continue to aid in clearing neuronal corpses (Figure S5). Notably, Spz3-depleted CG in the adult VNC often disappear completely rather than maintaining the globular morphology (Figure 6E’’-G’’), yet when compared to controls (Figure 6B’ and 6D’: astrocytes 2.94% ± 0.56% and SPG 4.64% ± 0.66%), Spz3^RNAi^ animals did not exhibit a significant increase in aberrant astrocyte or SPG infiltration (Figure 6E’,H and 6G’,J: astrocytes 3.48% ± 0.59% and SPG 6.11% ± 0.99%). In contrast, infiltration of EG in Spz3^RNAi^ animals (Figure 6F’,I, 20.34% ± 9.76%, 2.08-fold increase) significantly increased compared to control animals (Figure 6C’, 9.76% ± 1.24%). Moreover, the morphology of the infiltrating EG looked nearly identical to the wild-type CG lattice structure surrounding each neuronal cell body (Figure 6F’). This suggests that during metamorphosis and into early adult stages, the aberrant infiltration from larval astrocytes and SPG in CG Spz3^RNAi^ animals is removed, whereas EG processes dominate the cortex.

Given the differences between larval and adult infiltration, we sought to determine whether glial numbers might be altered in adults, so we again utilized anti-Repo (Figure S4A-D) and anti-Prospero (Figure S4G-J) antibodies while visualizing astrocytes (Figure S4A-B, G-H) or EG (Figure S4C-D, I-J) in the adult. There was no significant difference in the number of Repo+ neuropil-associated glia between control animals (Figure S4A,E, astrocytes: 620.54 ± 69.27; Figure 4B,F, EG: 478.42 ± 56.81) and CG Spz3^RNAi^ animals in each genetic driver condition (Figure S4E, astrocytes: 568.85 ± 55.85, Figure 4F, EG: 443.38 ± 43.69). Similarly, the number of Prospero^+^ nuclei is unchanged between control animals Figure S4K, astrocytes: 402.69 ± 44.16, L, EG: 320.5 ± 34.52) and Spz3^RNAi^ animals (Figure S4K, astrocytes: 408.5 ± 46.83, L, EG: 274.43 ± 24.58). These data reveal that even when CG are largely missing from the adult VNC and EG take over this region almost completely, the change in glial domains is not the result of altered numbers of the neighboring glial cells.

To determine if the neighboring glia continue to function in engulfing debris at this stage, we performed DCP-1 quantification in the same conditions tested in the larva. Like in larvae, control animals from astrocyte (Figure S5A,M, 3 ± 0.57), EG (Figure S5B,N, 13 ± 3.59), and SPG (Figure S5C,O, 15 ± 4.67) backgrounds were compared to conditions where (1) engulfment was blocked in the neighboring glia while CG remain unaltered (Figure S5D-F), (2) Spz3^RNAi^ is expressed in CG (Figure S5G-I) without directly altering neighboring glia, and (3) engulfment is blocked in the neighboring glia while Spz3^RNAi^ is expressed in CG (Figure S5J-L). While expressing Drpr^RNAi^ in astrocytes, there was a small increase in the amount of aberrant astrocyte infiltration into the cortex (Figure 6H), but no difference was observed in the EG (Figure 6I) or SPG (Figure 6J) backgrounds. There was no difference in the number of DCP1+ puncta in the control Draper knockdown condition in any of the glial driver backgrounds: astrocytes (Figure S5D,M, 9 ± 2.14), EG (Figure S5E,N, 17 ± 2.57), or SPG (Figure S5F,O, 20 ± 3.93). Likewise, we observed no change in DCP1+ puncta number with knockdown of Spz3 in CG in any driver background: astrocytes (Figure S5GM, 11 ± 2.91), EG (Figure S5H,N, 23 ± 5.81), or SPG (Figure S5I,O, 6 ± 1.67). In contrast to the L3 VNC, the number of DCP1+ nuclei in adult stages is quite low as developmental programmed cell death has largely ended by this point, so we would not expect the same build up of neuronal corpses. While we did observe slight differences in DCP1+ nuclei in the co-knockdown condition, these numbers remained quite low overall. Specifically, we observed a small increase in DCP1+ puncta in the astrocyte background (Figure S5K,M, from 11 ± 2.91 to 19 ± 5.52), a decrease in DCP1+ puncta in SPG labeled CNS (Figure S5L,O, from 6 ± 1.67 to 2 ± 0.59) and no change in the number of DCP1+ puncta in EG background (Figure S5K,N, 23 ± 5.81 to 14 ± 1.38). Together, these data suggest that glial interactions are dynamic, and their ability to take over glial territory and compensate for nearby glial dysfunction can change over the course of development or in different environmental contexts. Specifically, during metamorphosis, EG processes take over as the primary glial cell to infiltrate the cortex in the place of compromised CG. As programmed neuronal cell death is no longer as prevalent in adult stages, these infiltrating glia may no longer be performing a compensatory mechanism to clear neuronal debris, and may shift their effort to other supportive roles.

### Compensating glia continue to to perform their normal functions well

We have shown that astrocytes, EG, and SPG are able to compensate for missing cortex glial territory and function by aberrantly diverting their resources into the cortex to clear neuronal corpses in the larva, and EG take over in the adult. In order to ensure proper homeostasis, however, compensating glia would need to be able to perform their normal biological functions. Otherwise, compensation for neighboring glial dysfunction would be detrimental even to regions outside of the initial dysfunction. To determine if astrocytes, EG, and SPG are all still capable of performing their normal functions while compensating for impaired CG function, we investigated well-characterized functions of each subtype: astrocytic clearance of synaptic debris during early pupal formation,^59,62^ EG clearance of axonal debris after injury in the adult,^1^ and the ability of larval SPG to maintain the integrity of the BBB.^34^

To investigate astrocytic function at a time when they still utilize a portion of their cellular resources to extend processes from the neuropil into the cortex, we focused on their role in clearing synaptic debris at the onset of pupal formation. Sixteen VNC peptidergic Corazonin (vCrz)^+^ neurons undergo apoptosis during early metamorphosis between 0 and 6 hours after pupal formation (h APF), at which point their neurites and cell bodies are eliminated by astrocytes and CG, respectively.^59,62^ To fully track the ability of astrocytes to clear vCrz neuronal debris and ensure that initial development of these neurons was not perturbed, we first visualized vCrz neurons using an anti-Crz antibody at L3 and quantified the number of vCrz cell bodies and neurite volume. In both control (Figure 7A) and Spz3^RNAi^ (Figure 7C) animals, we observed all 16 cell bodies and neurites, indicating that there is no difference in initial vCrz neuron development between the two conditions (Figure 7E). By 6h APF in control animals, most vCrz cell bodies are cleared by CG (Figure 7B,D,E; 4.75 ± 0.85) and the neurite debris cleared by astrocytes (Figure 7B,D,F; 3927 ± 641.40 um^3^). In Spz3^RNAi^ animals at 6h APF, more vCrz+ neuronal somas remained, demonstrating the decreased ability of globular CG to clear vCrz neuron cell bodies (Figure 7D- E; 8.18 ± 0.99; 1.72-fold increase) since they no longer surround the majority of these somas; however, even with their aberrant processes extending into the cortex, astrocytes in Spz3^RNAi^ animals are still able to clear out the neurite debris on par with control animals (Figure 7D,F; 3595 ± 831.54 um^3^).

**Figure 7.**
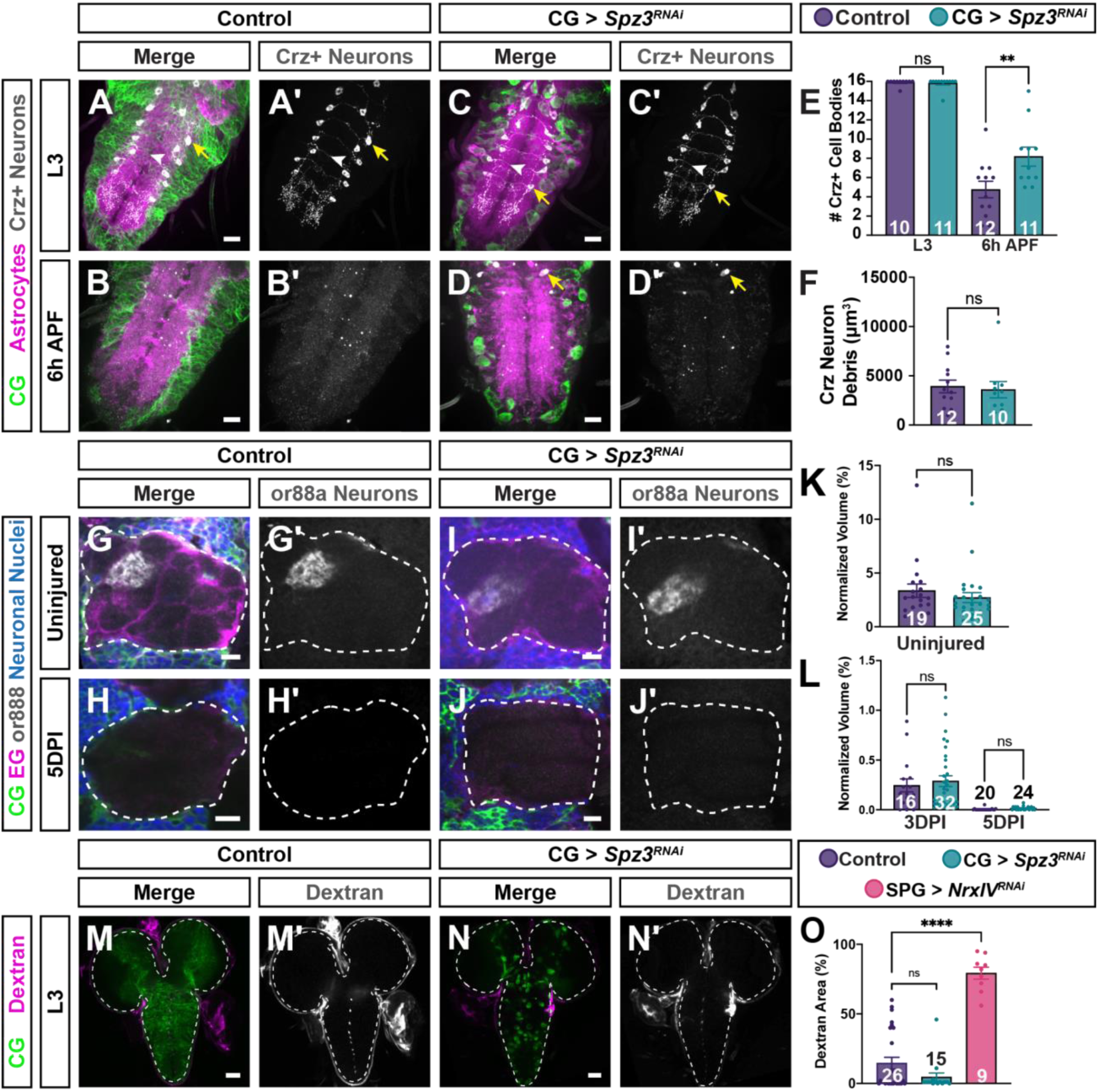
Glia that compensate for neighboring glial dysfunction are still capable of performing their normal functions. (**A-B**) Control VNCs with CG (green), astrocytes (magenta), and Crz neurons (anti-Crz, gray) show normal Crz+ neuronal cell bodies and projections at L3 (A), but by 6 hours after pupal formation (h APF) the cell bodies are mostly cleared by CG and the projections are cleared by astrocytes (B). (**C-D**) When CG expressing RNAi against Spz3 exhibit disrupted CG morphology, more Crz+ neuron cell bodies remain, while the projections are still cleared. Scale bars = 10 µm. White arrowheads denote Crz+ neuronal projections while the yellow arrowheads denote the Crz+ neuronal cell bodies. (**E**) Quantification of the number of Crz+ neuronal cell bodies at L3 and 6h APF and (**F**) Crz neurite debris remaining at 6h APF. (**G-H**) Control brains with CG (green), EG (magenta), neuronal nuclei (blue), and or88a-CD2 (anti-ratCD2, gray) show or88a+ neuronal projections in the antenna lobe in uninjured animals (G), but these projections are cleared by EG following injury partially at 3 days post injury (DPI) and fully by 5 DPI (H). (**I-J**) When CG morphology is disrupted with Spz3 RNAi, or88a+ neurons are still cleared by EG. Scale bars = 10 µm. (**K-L**) Quantification of the or88a+ volume normalized to the volume of the antennal lobe in uninjured control and Spz3 knockdown animals (K) and at 3 and 5 DPI (L). (**M**) Control brains with CG (green) exposed to 70 kDa Dextran (magenta) successfully block Dextran from the L3 CNS. (**N**) When CG express Spz3^RNAi^, Dextran is still excluded from the L3 CNS, suggesting an intact BBB and functioning SPG. (**O**) Quantification of the area of Dextran within the L3 CNS. N = numbers of animals, denoted in each corresponding bar. (n.s.) Not significant. (*) *P* < 0.5, (**) *P* < 0.01, (***) *P* < 0.001, (****) *P* < 0.0001. (E, F, K) Unpaired t-test, (L,O) One-Way ANOVA with multiple comparisons.

To assess EG function, we focused on their ability to engulf severed olfactory receptor neuron (ORN) axons. ORNs located within the antenna and maxillary palps send projections to one of over 50 distinct glomeruli within the antennal lobes of the brain.^63^ As the pattern of the projections are spatially invariant, they can easily be tracked during development and degeneration. When these neurons are severed via antennal removal, they undergo Wallerian degeneration^1^ and the axonal debris is subsequently cleared by EG with stereotyped timing: partial clearance by 3 days post injury (DPI) and full clearance by 5 DPI. Therefore, post-injury axon clearance can be used to determine whether the EG can retain their normal functions even when they infiltrate the cortex after impaired CG function. Utilizing endogenously-expressed *CD2-*tagged Or88a (Or88a-CD2)^64^ in conjunction with an anti-CD2 antibody, we visualized a subset of olfactory neuron projections into a singular glomerulus of the antennal lobe and quantified the volume of or88a debris, normalizing to total antennal lobe volume. Animals in control (Figure 7G,K; 3.35% ± 0.63%) and Spz3^RNAi^ (Figure 7I,K; 2.71% ± 0.45%) conditions have similar volumes of Or88a axons present prior to injury. After injury is induced by manual antennal ablation, EG partially clear the debris by 3 days post injury (DPI) and completely by 5 DPI in both control (Figure 7H,L; 3DPI: 0.24% ± 0.06%, 5DPI: 0.00% ± 0.00%) and when EG infiltrate into the cortex in CG Spz3^RNAi^ conditions (Figure 7J-L; 3DPI: 0.29% ± 0.05%, 5DPI: 0.01% ± 0.00%). These data indicate that EG are still able to perform one of their major functions just as well even with a portion of their cellular resources diverted to the cortex.

Septate junctions in SPG make them integral members of the BBB, blocking small molecule diffusion into the CNS.^32^ To determine whether SPG that aberrantly invade the cortex are still sufficient in their role in blocking small particle diffusion into the CNS, we exposed L3 CNS tissue to 0.6 mg/ml of 70 kDa Dextran (Figure S6A) conjugated to Tetramethylrhodamine dye in control animals (Figure 7M,O), those with CG expressing Spz3^RNAi^ (Figure 7N,O), and positive controls where Neurexin IV (NrxIV) was knocked down in SPG (Figure 7O, Supp Figure 6B). NrxIV is a membrane protein essential for proper septate junction formation, and its loss from SPG is known to disrupt the BBB.^65^ When NrxIV^RNAi^ is expressed specifically in the SPG, Dextran readily passed through the BBB, as measured by intensity (Figure S6B-C; 636.34 ± 82.65) and area (Figure 7O; 79% ± 4.29%) of Dextran found within the CNS at L3. In the control L3 CNS (Figure 7M,O; Dextran Intensity: 301.22 ± 51.46, Dextran Area: 15% ± 4.25%), SPG are efficient at keeping the Dextran dye outside the CNS. When CG express Spz3^RNAi^ and SPG infiltrate into the cortex, there is no statistically significant difference in Dextran in the CNS (Figure 7N,O; Dextran Intensity: 173.07 ± 15.02, Dextran Area: 5% ± 3.04%), demonstrating that infiltrating SPG maintain their ability to block diffusion and the BBB remains intact. Overall, these data demonstrate that in a context where one glial subtype can no longer perform its own functions, surrounding glia divert a portion of their resources to infiltrate beyond their typical boundaries and perform compensatory actions, multitasking to take on new roles while still performing their normal biological functions.

## Discussion

Within the CNS there is extensive glial-glial communication needed to ensure proper interactions with and support to the neurons. However, detailed studies of glial-glial physiological functions *in vivo* have been hindered by the complexity of both neuronal and glial architecture in vertebrate systems. Because of this, while tiling is known to occur between mammalian glia within a single subtype,^12,13,21–24^ the field is only beginning to understand the full extent of the mechanisms that regulate how glial cells interact and communicate across subtypes. It is therefore difficult to gain a clear understanding as to how the dysfunction of one type affects healthy neighboring glia amidst the complexity of the mammalian nervous system, or how these otherwise healthy glial cells react and compensate for dysfunctional glia. In addition to the complexity, the lack of genetic tools to easily target and manipulate multiple specific glial subtypes simultaneously in mammalian models has precluded the advancement of this field. In this study, we investigated the dynamics between altered *Drosophila* CG and their glial neighbors to overcome these limitations due to the advantageous CNS organization, versatile genetic tools, and functional conservation between *Drosophila* and mammalian glia. Our data demonstrate that in a context where one glial subset is morphologically and functionally disrupted, all closely interacting neighboring glial cell types invade the previously occupied territory. This aberrant outgrowth appears not to be the result of increased glial proliferation or migration, but instead enhanced aberrant process extension into the cortex, functioning at least in part as a compensatory mechanism to clear neuronal cell body corpses. Finally, we show that even when astrocytes, EG, and SPG compensate for the lack of CG function, they are still capable of performing their normal biological roles, highlighting the ability of these cells to multitask in order to preserve nervous system function.

### Glia respond to impaired glial neighbors by taking over their territory and compensating for lost functions

Nearly every type of glial cell tiles to form domain boundaries with neighboring cells of the same glial subtype,^12,21,22,24,38,66–68^ but *Drosophila* allows for a unique ability to understand how different glial cell types interact across compartmentalized domains. Previously, we have shown that upon genetic ablation of CG or even just simply their morphological disruption, astrocytes extend processes to infiltrate into the previously CG territory.^36^ These data align with a subsequent study in mice, in which microglia and astrocytes display a similar relationship where microglia are the primary glial cell to engulf neuronal cell body debris, yet astrocytes take over for this action upon the loss of microglial.^39^ Whether this compensation occurs at the expense of the cell’s own functionality was not clear. Additionally, in an *in vivo* system, glial cells form complex interactions among many glial subtypes. Here, we demonstrate that all of the glial cells that interact with CG are capable of responding to their cellular dysfunction by growing into the cortex to help replace lost functionality. This is highlighted by the fact that early ablation of CG is known to result in animal lethality in early developmental stages.^36,69^ As opposed to complete loss of CG all at once, the gradual loss of CG territory and function likely allow the neighboring glia time to infiltrate into the cortex and compensate for CG functions. However, our results demonstrate that not all glial cells have the same capacity to compensate for the lost function, and the ability to do so can change under different contexts such as in distinct developmental stages and throughout lifespan. For example, after the prolonged depletion of Spz3 specifically in CG, the morphological phenotype worsens from globular structures in the larval CNS to being largely absent in the adult VNC. At these later stages, not only are EG the predominant glial cell to take over interactions with neuronal somas, but they could also have a direct response in clearing the previously occupying CG cells such as microglial and astrocyte engulfment of oligodendrocyte myelin debris in vertebrates.^70^ The full extent of this compensatory capacity, and whether different glial cells are more capable of recovering certain functions than others is currently unclear, underlining the need within the the field to gain a better understanding of how different neighboring glial cells respond to neurons and other impaired glia throughout lifespan or in different disease contexts. In diseases where glial cells are genetically or functionally transformed, such as in glioma^71^ or demyelinating oligodendrocytes in Multiple Sclerosis,^72^ understanding how neighboring glia react at different time points in disease could shed light on new aspects of the underlying biology or novel ways to treat each.

The current study focused on engulfment of neuronal debris, a well-known role in nearly all glial subtypes;^1,4,39,59,73–86^ however, phagocytosis is only one of the many functions carried out by glial cells. CG not only clear neuronal corpses throughout development and disease,^4^ but also support neurons by providing neuronal metabolic support^37^ and ion buffering,^87^ as well as regulate neuroblast proliferation.^88,89^ How these supportive roles are affected by cortex glial loss, and the extent to which astrocytes, EG, and SPG can compensate for these functions is currently unknown. Moreover, whether glial cells in any system have the capacity to recapitulate all lost functions equally, in which contexts each cell optimally performs each function, and how many new roles each cell can take on before its own functions are impaired are intriguing areas of investigation that have only begun to be explored.

### Infiltration of aberrant glial processes is not linked to proliferation

CG exhibit self-tiling and compensatory growth behavior to the extent that when a single CG is ablated, the surrounding CG will grow into the domain of the dying CG cell to replace the glial wrapping of nearby neuronal somas.^36^ This tiling behavior and ability to grow to fill neighboring territory is lost when all CG simultaneously transform into globular cells that no longer proliferate. Given the dramatic increase in the number of CG nuclei that occurs in control animals during the L2-L3 CNS expansion, it was unclear whether this proliferation was necessary for CG growth and full territorial coverage. Our current work demonstrates that an increase in CG proliferation is not required to maintain full coverage of CG processes throughout the cortex, nor can it rescue the globular morphology and territorial loss. Although CG remain capable of encapsulating old and new neuronal cell bodies when their proliferation is blocked, it is unclear whether wrapping by a stagnant number of cortex glial cells can provide optimal function at the glial-somal interactions; however, the fact that astrocytes, EG, and SPG cells are able to increase their outgrowth to take on novel functions while preserving their own suggests that these expanded cortex glial cells might be just as capable. Accordingly, we found that the number of astrocytes and EG do not increase in order to compensate for the loss of cortex glial function. Recent work focusing on astrocyte-ensheathing glial interactions revealed that after ablation of EG, astrocytes increase in number.^10^ Additionally, genetic ablation of neurons has been shown to lead to an increase in Prospero+ glia in the embryo,^90^ and stab injury in the larval CNS induces proliferation of Prospero+ neuropil-associated glia.^91^ The lack of proliferation of the neighboring glia that react to CG dysfunction therefore implies that this response is separate from an injury response, suggesting that alternative signaling underlies the aberrant growth and functional compensation.

### Genetic manipulation of neuronal cell death impacts aberrant glial growth into the cortex

Although multiple mechanisms of cell death exist, programmed cell death through apoptotic pathways plays an important role in many aspects of development and disease.^92^ Indeed, we know that CG play an major role in clearing apoptotic neuronal corpses throughout embryonic, larval, and pupal stages, and a failure to do so can result in neurodegeneration in the adult.^4,50,93^ Specifically, dying neurons have been shown to secrete a separate Spz family member, Spz5, to prime CG for phagocytosis through activation of Toll-6 and its downstream signaling factors, eventually resulting in the upregulation of Draper for phagocytosis and corpse removal.^4^ We therefore hypothesized that upon the loss of cortex glial interactions, Spz5 or other signals from these dying neurons could instead reach neighboring glia to prime them, aiding in the clearance of dying neuronal cell bodies. By blocking effector caspases like DCP-1 and DrICE with the genetic expression of the baculovirus p35 gene, we initially expected to reduce cell death and potentially the extent of aberrant infiltration. However, the expression of p35 throughout development, when these apoptotic pathways naturally turn on in cells programmed to die, could ultimately lead to an accumulation of “undead” cells.^57^ In particular, not only does the initiator caspase signaling upstream of the effector caspases still occur, but these undead cells have also been shown to cause non-autonomous effects on proliferation and survival of surrounding cells via enhanced Wnt, TGF-β, Hedgehog, and EGFR signaling, among others.^94,95^ CG rise a glial niche for both quiescent and proliferating neuroblasts and have been documented to signal bi- directionally with these stem cells;^88,89,96^ therefore, changes in neuroblast proliferation could alter their progeny, the remaining CG, or even other glial cells attempting to compensate for cortex glial functions.

The 3xP3-RFP transgene used in our neuronal p35 expression experiments visualizes SPG and EG, but does not readily label astrocytes (Figure S3), therefore our data likely underestimate the changes that occur due to infiltration of all neighboring subtypes; however, even with that caveat, our data demonstrate that the expression of p35 leads to increased neighboring glial infiltration. It is currently unclear which specific cues (e.g. Spz5, Wg, Dpp, or others) signal to these neighboring glial processes, and whether the cues result from a call from unwrapped neurons asking for more glial engulfment, enhanced neuronal support from surviving neurons, or even lost communication that arises from CG themselves. This work highlights the need to better understand the broad signaling mechanisms that regulate glial-glial territories, how those territories relate to function and neuronal support, the extent of the plasticity in their functional compensation, and how and which neuronal cues play a role in turn.

## Acknowledgements

We thank all members of the Coutinho-Budd laboratory for helpful advice and discussions. We are grateful to the Vienna Drosophila RNAi Center and the Bloomington Stock Center for providing various fly stocks, Dr. Hermann Steller for sharing the QUAS-p35 *Drosophila* stock, and to Dr. Marc Freeman for providing the anti-Crz antibody. We thank Colleen McLaughlin, Brandon Holmes, Megan Corty, Sarah Kucenas, and Bettina Winckler for providing helpful feedback on the manuscript. This work was supported by the National Institutes of Neurological Disorders and Stroke (R01NS121101 to JCB).

## Author Contributions

Experiments performed in larval animals were completed by G.S., K.K., A.N., H.K., K.M., A.M., G.R., H.B., L.A., H.M, and J.C.B. Experiments performed in adult animals were completed by A.B. and H.K. qPCR was performed by T.W. J.C.B conceived of and supervised this project. Manuscript was written by A.B and J.C.B.

## Materials and Methods

### Fly Strains

Unless otherwise noted, *Drosophila melanogaster* crosses were set on Molasses Food (Archon Scientific) at 25℃ and then progeny were moved to 29℃ until larvae reached L3. For adults, animals were raised at 25℃ until adult flies eclosed, at which time the animals were housed at 29℃. The following previously made *D. melanogaster* transgenes were used in this study: *CtxGlia-SplitGal4,*^36^ *GMR25H07-LexA* (BDSC#52711)*, GMR54C07-LexA* (BDSC#61562)*, GMR56F03-Gal4* (BDSC#39157)*, UAS-CD8::GFP,*^97^ *LexAop-rCD2::GFP* (BDSC#66544)*, GMR54C07-Gal4* (BDSC#50472)*, Elav-QF2(PxP3-RFP)* (BDSC#66466)*, lexAop-CD8-GFP* (BDSC#32203)*, UAS-mCherry.nls* (BDSC#38424)*, UAS-LacZ.nls,*^98^ *UAS-Spz3RNAi^102871^* (VDRC#102871)*, LexAop-Spz3RNAi^102871,^*^36^ *UAS-stg.HA* (BDSC#56563)*, UAS-CycARNAi^103595^* (VDRC#103595)*, UAS-StgRNAi^17760^* (VDRC#17760)*, UAS-StgRNAi^TRiP.HMS00146^* (BDSC#34831)*, LexaOp2-DraperRNAi^M2^,*^36^ *pWIZ-UAS-DraperRNAi[7B],*^1^ *UAS-NrxIVRNAi^108128^* (VDRC#108128), *QUAS-p35,*^99^ *QUAS-rpr* (BDSC#92782), and *or88a-CD2* (BDSC#23298). Genotypes used in each figure are specified in Supplemental Table 1.

### Immunohistochemistry and Imaging

L3 CNSs were dissected in 1x PBS, fixed in ice-cold 4% Formaldehyde (PBS + 4% Formaldehyde) for 25 minutes before washing with 0.1% PTX (PBS containing 0.1% Triton-X) for 5 minutes (3x). Subsequently, tissue was stained overnight at 4℃ with antibodies in 0.1% PTX. For adult dissections, adult CNSs were dissected 5 days post eclosion (DPE) in 0.3% PTX (PBS + 0.3% Triton-X), fixed in ice-cold 1% Formaldehyde (PBS + 1% Formaldehyde) for 2 hours before washing with 0.3% PTX for 5 minutes (x3). To optimize tissue quality, tissue was fixed within 20 minutes of tissue harvesting. Subsequently, tissue was stained over 2 nights at 4℃ with antibodies in 0.3% PTX.

The following primary antibodies were used: chicken anti-GFP (1:1000; Antibodies Inc), rat anti-mCherry (1:1000, Invitrogen); rabbit anti-DsRed (1:1000; Clontech) rat anti-Elav (1:100; Developmental Studies Hybridoma Bank [DSHB], 7E8A10), mouse anti-Elav (1:100, DSHB, 9F8A9), Anti-β-Galactosidase (1:1000; Promega, Z3781), mouse anti-Repo (1:5; DSHB, 8D12), mouse anti-Prospero (1:100; DSHB, MR1A), rabbit anti-DCP1 (1:100; Cell Signaling, 9578S), rabbit anti-corazonin (1:500, Freeman Lab), mouse anti-nc82 (1:20; DSHB, nc82), mouse anti- ratCD2 (1:200; BioRad, MCA154GA), rabbit anti-p35 (1:1000, Fisher Scientific, NB10056153T). Primary antibodies were detected with the appropriate goat or donkey secondary antibodies conjugated to Dylight405 (rabbit [711-475-152], mouse [715-475-151], and rat [712-475-153]), Dylight488 (chicken [103-005-155]), Cy3 (rabbit [711-165-152], mouse [715-165-151], and rat [712-175-153]), or Cy5 (rabbit [711-175-152), mouse [715-175-151], and rat [712-175-150]) from Jackson ImmunoResearch. Samples were mounted in VectaShield reagent (Vector Laboratories, H-1000) and imaged on an Innovative Imaging Innovations (3I) spinning-disc confocal microscope equipped with a Yokogawa CSX-X1 scan head.

### Growth Restriction Assay

Eggs were laid in vials of Molasses Food (Archon Scientific) and placed at 25 °C. L1 animals were removed from the surface of the food, separated by condition, and transferred onto 35 mm petri dishes containing 20% sucrose/3% agar in H_2_O. Each dish was placed inside a 10 cm square petri dish with a wet Kimwipe to serve as a humidity chamber. Animals were raised on the sucrose agar at 25 degrees for an additional 5 days, during which time they remained approximately the same size. Viable animals from each genotype were then removed, the CNS was dissected, fixed in 4% formaldehyde in PBS for 20 minutes at room temperature, washed, and stained via immunohistochemistry.

### RNAi Isolation and qPCR

Whole adult flies 5 DPE were flash frozen in dry ice prior to head removal. RNA was isolated according to manufacturer’s instructions with the Qiagen RNeasy Mini kit. After RNA isolation, RNA was eluted in 30ul ddH2O. Input for cDNA synthesis was 1ug RNA per reaction. Synthesized cDNA was diluted 1:4 in ddH2O, and 2ul cDNA was used per reaction. Primer stock concentrations were 100uM, primer final concentrations in the reactions were 500nM. cDNA synthesis was performed according to manufacturer’s instructions with the BioRad iScript cDNA Synthesis kit. cDNA was used in the qPCR reaction using a iTaq Universal SYBR Green Supermix (BioRad). The qPCR was performed on BioRad CFX Opus 384 and analyzed with CFX Maestro software. Each qPCR reaction was run in triplicate. CT values of RpL32 were measured as a quality control housekeeping gene, and relative expression of Rpr was determined by calculating 2^-ΔCt, using RpL32 expression levels for normalization. The following primers were used:

rpr Fwd 5’- GAGTCACAGTGGAGATTCCT -3’

rpr Rev 5’- GATGGCTTGCGATATTTGCC -3’

RpL32 Fwd 5’- ATGCTAAGCTGTCGCACAAA -3’

RpL32 Rev 5’- AACCGATGTTGGGCATCAGA -3’

### Antennal Ablation

Antennal injury performed using a previously described protocol (Vosshall et al., 2000). Briefly, both antennae were removed with forceps from anesthetized 3DPE adult flies. Animals were allowed to recover in fresh molasses food vials and aged for 5 additional days at 25°C before projection patterns were analyzed via immunohistochemistry.

### Dextran

Larva: Third instar larvae (L3s) were collected into one well of a micro spot plate containing PBS. Forceps were used to remove the posterior-most portion of the larva and invert the larval body. Once the larvae had been inverted, the PBS was replaced with 70kDa Dextran (Dextran, Texas Red D1830, Fisher Scientific) in PBS at a concentration of 0.58 mg/ml–the effective concentration that showed a significant difference between infiltration into the negative and positive controls (Figure S6A). The tissue was incubated on an orbital shaker at low speed for 40 minutes. VNCs were then washed with PBS and subsequently fixed in 4% formaldehyde in PBS for 20 minutes, washed twice with PBS for 5 minutes and then twice with 0.1% PTX for 10 minutes at a time. The CNS was then dissected, placed in Vectashield and mounted onto a glass slide for imaging.

### Analysis and Statistics

Imaris and ImageJ softwares were used for all quantifications. For infiltration assessment, the volume of the cortex was measured in Imaris by generating a surface based on the area containing neuronal cell bodies throughout the Z-series. The volume of glial cell types were measured using a surface generated based on ImageJ LabKit segmentation. The volume of aberrant infiltration into the cortex by neighboring glial subtypes was quantified by selecting for the overlap between these two surfaces. For experiments involving 3xP3-RFP, which also labels cortex glia, the cortex glial channel was subtracted from the total glial channel to include quantification of only invading neighboring glial cells. For cell death, the number of dying neurons was visualized using anti-DCP1 expression, and DPC-1+ nuclei were identified using ImageJ Labkit Segmentation and quantified in Imaris. To quantify neuropil-associated glial cell number, the number of glial cell nuclei was visualized using anti-Repo expression or anti-Prospero expression and puncta were manually identified and quantified in Imaris.

Corazonin cell bodies were visualized using anti-corazonin expression and quantified in Imaris. Corazonin synaptic debris was visualized using anti-corazonin expression and a Imaris surface was created by thresholding the absolute intensity of that channel. The surface was secondarily filtered with the minimum area was 0.00 µm^2^ and the maximum was 200 µm^2^ to allow for the inclusion of synaptic debris while excluding the larger cell bodies. Antennal lobe volume was measured by generating a surface of the antennal lobe volume throughout the Z-series. The volume of Or88a+ neuronal projections was then measured by generating a surface of the or88a- CD2+ channel using ImageJ Labkit segmentation. The volume of Or88a+ neuronal projections from the left and right antennal lobes were averaged together for each animal, and Or88a volume was normalized to the volume of the antennal lobe for each animal.

For Dextran infiltration, the mean intensity was quantified in Imaris by generating a surface of the Dextran channel within the neuropil half-way through the CNS. The area of Dextran infiltration was generated by creating a surface for the absolute intensity of the Dextran channel, with a minimum intensity set to 600 ms. The area was divided by the total neuropil area to generate a percentage of Dextran infiltration.

Error bars in all graphs indicate ± SEM. Statistical significance was calculated with Graphpad Prism 10 software by statistical analysis with an ANOVA with the appropriate post hoc test for comparing more than two sets of data, or unpaired t-test when only two conditions were present.

## Supplemental Titles and Legends

**Figure S1.**
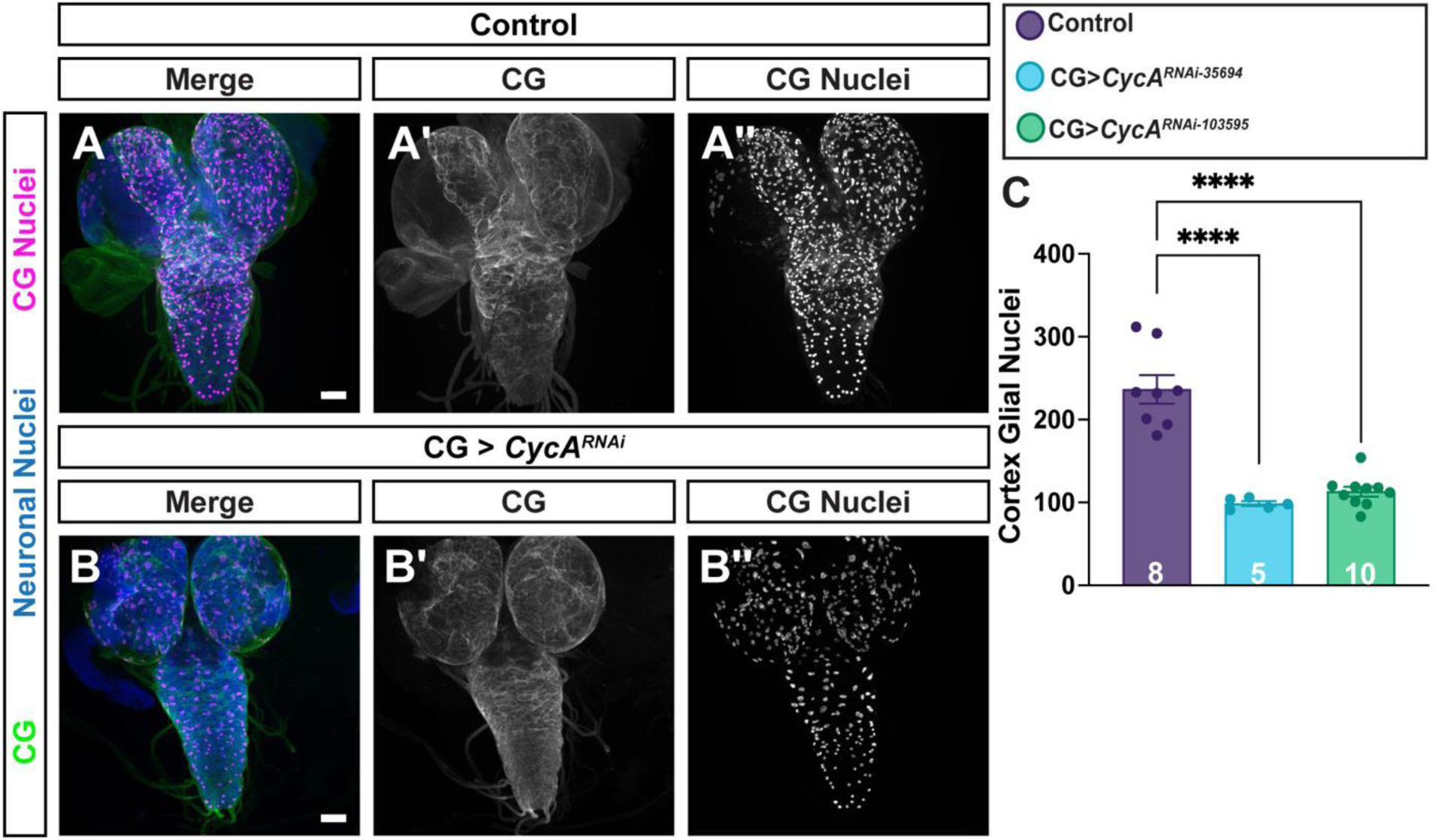
CG proliferation is not required to maintain morphology, related to. Figure 3. (**A**) Control animals with CG (green, grayscale A’), neuronal cell bodies (anti-elav; blue), CG nuclei (UAS-mCherry.nls, magenta, grayscale A”). (**B**) CG expressing CycA^RNAi^ exhibit no alteration to CG morphology (grayscale, B’), but a significant decrease in the number of CG nuclei (grayscale, B”). Scale bars = 50 µm. (**C**) Quantification of CG nuclei in the VNC. N = numbers of animals, denoted in each corresponding bar. (n.s.) Not significant. (*) *P* < 0.5, (**) *P* < 0.01, (***) *P* < 0.001, (****) *P* < 0.0001. One-Way Anova, with multiple comparisons.

**Figure S2.**
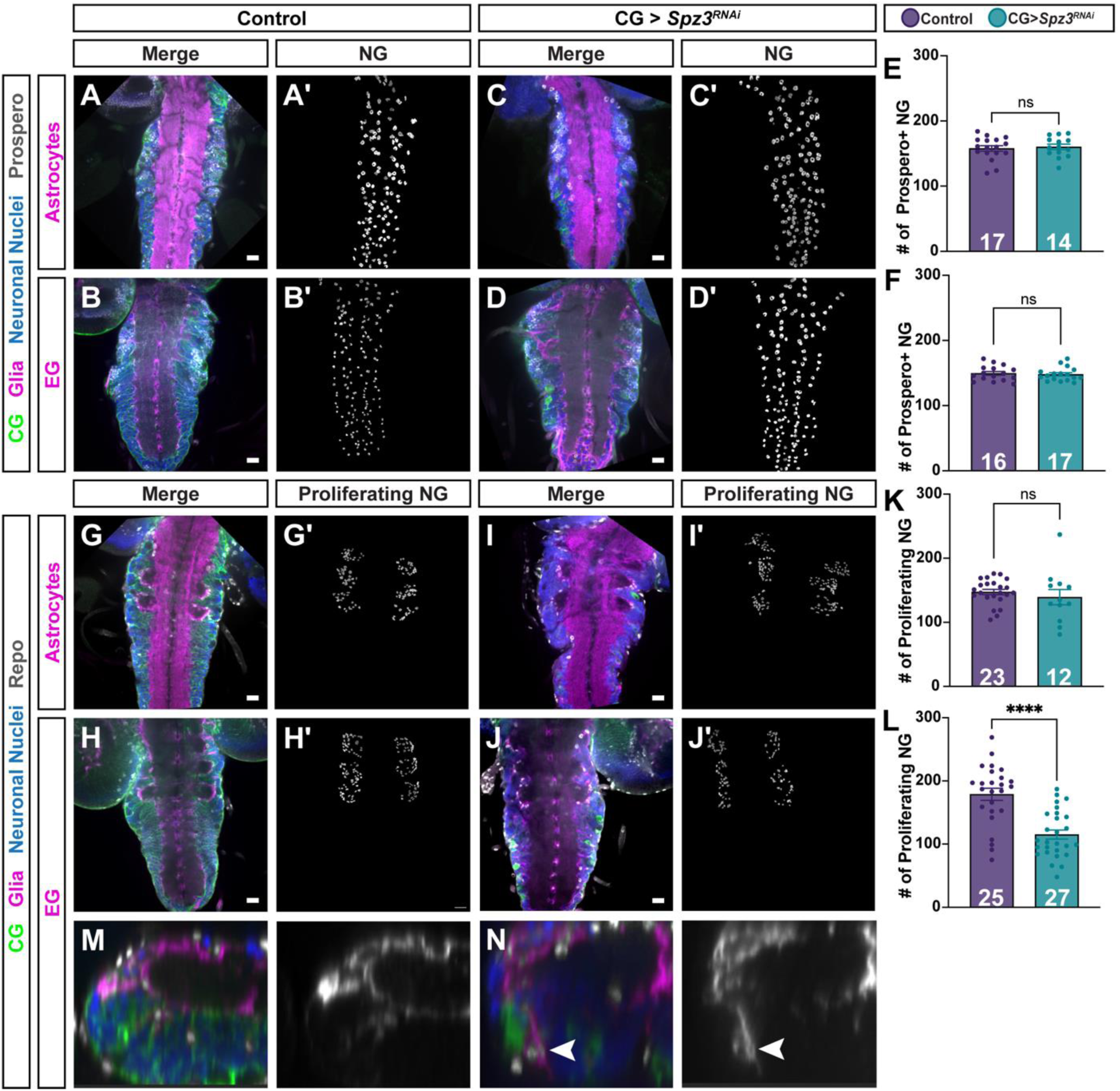
Neuropil-associated glial numbers do not change in response to disrupted CG but the number of proliferating glia is altered, related to. Figure 3. (**A**) Spz3 knockdown animals with CG processes (green), glia nuclei (gray) and astrocytes (magenta) depict normal CG throughout the VNC when raised on 20% sucrose agar. Grayscale image of selected neuropil-associated glia only (A’). (**B-D**) Quantification of Prospero+ neuropil-associated glia (NG) nuclei. (**E-F**) Control animals with CG (green), neuronal cell bodies (anti-elav; blue), astrocyte nuclei (anti-prospero; gray) with labeled astrocyte (E) and EG (F) cells in magenta. (**G-J**) CG expressing Spz3^RNAi^ do not exhibit a statistically significant difference in the number of neighboring Prospero+ NG nuclei. Grayscale images of NG only (E’-H’), quantified in I-J. (**K-L**) Control animals with CG (green), neuronal cell bodies (anti-elav; blue), proliferating glial precurosor nuclei (anti-repo; gray) with labeled astrocytes (K) or EG (L) in magenta. (**M-P**) Animals with CG expressing Spz3^RNAi^ exhibit no change in proliferating NG glia in the astrocyte driver background, but reduced number in the EG driver condition (N). Grayscale images of selected proliferating NG only (K’-N’), quantified in O-P. Scale bars = 20 µm. N = numbers of animals, denoted in each corresponding bar. (n.s.) Not significant. (*) *P* < 0.5, (**) *P* < 0.01, (***) *P* < 0.001, (****) *P* < 0.0001. Unpaired t-test. (**Q-R**) Orthogonal view of control (Q) and Spz3RNAi (R) L3 VNC with CG (green), neuronal cell bodies (anti-elav, blue), glial nuclei (anti-Repo, gray), and EG (magenta). Grayscale image of EG channel only (Q’-R). White arrowhead highlights a migrating EG cell body into the cortex.

**Figure S3.**
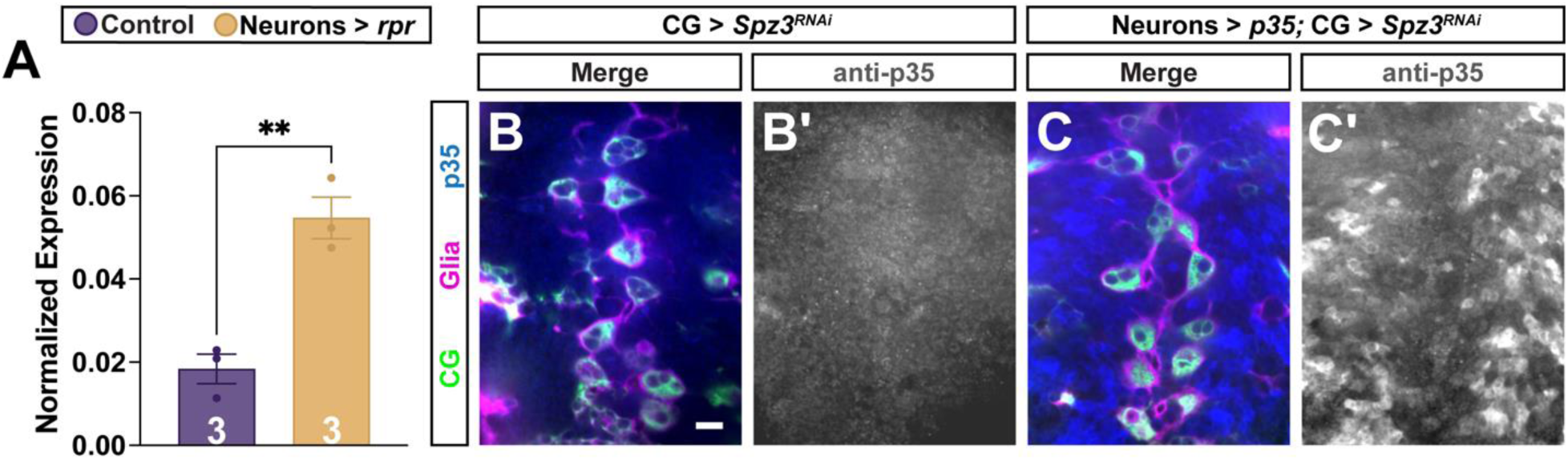
Overexpression constructs of rpr and p35 in neurons results in increased expression, related to. Figure 5. (**A**) Quantification of rpr mRNA by qPCR demonstrates increased expression of rpr in animals driving neuronal *rpr* expression. N = numbers of samples, each containing 50 brains, denoted in each corresponding bar. (**) *P* < 0.01. Unpaired t-test. (**B-C**) Anti-p35 antibody demonstrates no expression of p35 in control animals (B), and successful expression of p35 in animals with neuronally-driven p35 (C). CG (green), glia (3xP3- RFP, magenta), neuronal nuclei (anti-Elav, blue), and p35 (anti-p35, gray, B’,C’) Scale bars = 20 µm.

**Figure S4.**
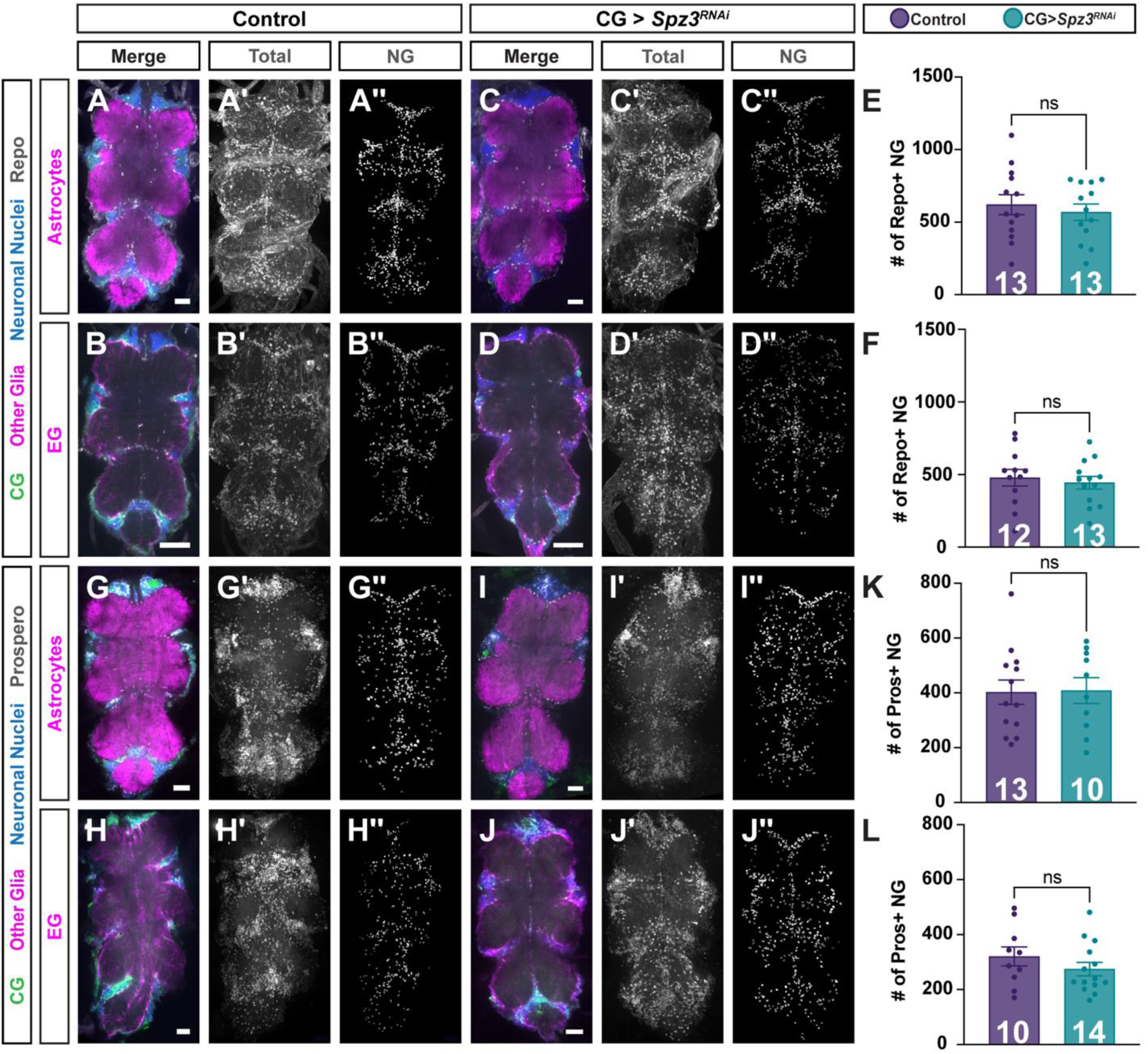
Number of neuropil-associated glia do not change in response to disrupted CG in adults, related to. Figure 6. (**A-B**) Control adults show normal CG (green), neuronal cell bodies (anti-elav, blue), glial nuclei (anti-Repo, gray) with neuropil-associated astrocytes (A) and EG (B) in magenta. (**C-F**) CG expressing RNAi against Spz3 does not change the number of Repo+ neuropil-associated glia in the astrocyte (C) or EG (D) driver backgrounds, quantified in E and F, respectively. (**G-L**) Control adults show normal CG (green), neuronal cell bodies (anti-elav, blue), with Prospero+ glial nuclei (anti-Prospero, gray) and neuropil-associated astrocytes (G) and EG (H) in magenta. (**I-J**) CG expression of RNAi against Spz3 does not change the number of neuropil-associated glia in the astrocyte (I) or EG background (J), quantified in K and L, respectively. Grayscale images of the Repo channel (A’-B’, G’-J’) and selected neuropil-associated glia (A’’-B’’, G’’-J’’) alone. Scale bar = 100 µm. N = numbers of animals, denoted in each corresponding bar. (n.s.) Not significant. Unpaired t-test.

**Figure S5.**
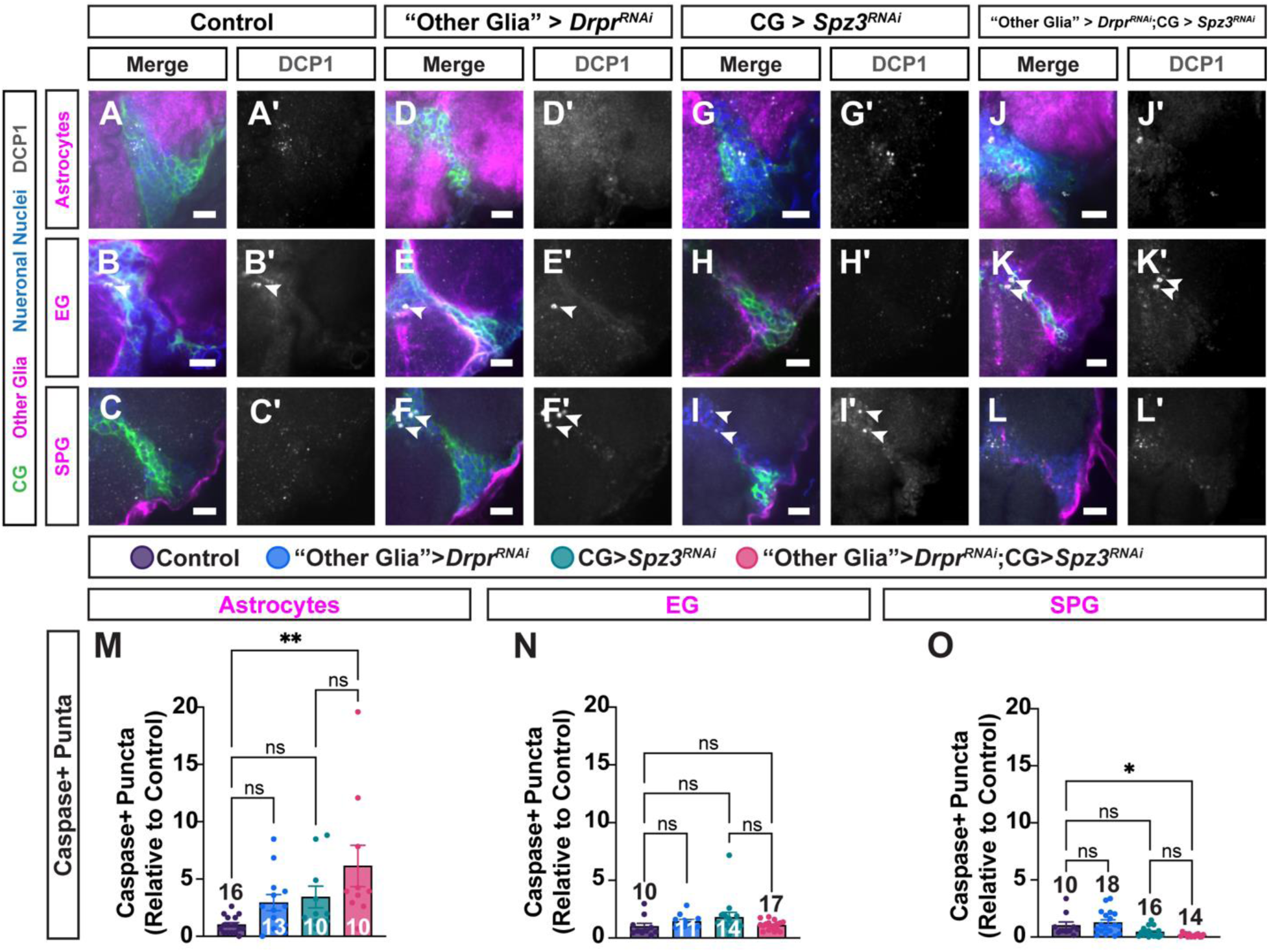
Infiltrating glia do not clear DCP1+ corpses in adult VNC, related to. Figure 6. (**A-L**) DCP-1 staining in adult VNCs with CG (green), astrocytes, EG, or SPG (magenta), neuronal nuclei (anti-Elav, blue) and DCP-1+ nuclei (anti-DCP1, gray). (**A-C**) Control brains exhibit minimal numbers of DCP1+ puncta. (**D-F**) Loss of Drpr in each non-CG subtype with wild-type CG results in minimal numbers of DCP1+ puncta. (**G-I**) CG expressing RNAi against Spz3 results in disrupted CG morphology, but no increase in DCP-1+ cells. (**J-L**) Animals with both Spz3 knockdown in CG and Drpr knockdown in either (J) astrocytes, (K) EG, or (L) SPG do not mimic the universal increase in DCP-1+ cells seen in this condition at L3. Grayscale images of the DCP1 channel alone (A’-L’). Scale bars = 50 µm. White arrowheads denote DCP-1+ puncta. (**M-O**) Quantification of the number of DCP1+ puncta. N = numbers of animals, denoted in each corresponding bar. (n.s.) Not significant. (*) *P* < 0.5, (**) *P* < 0.01. Two-Way ANOVA with multiple comparisons.

**Figure S6.**
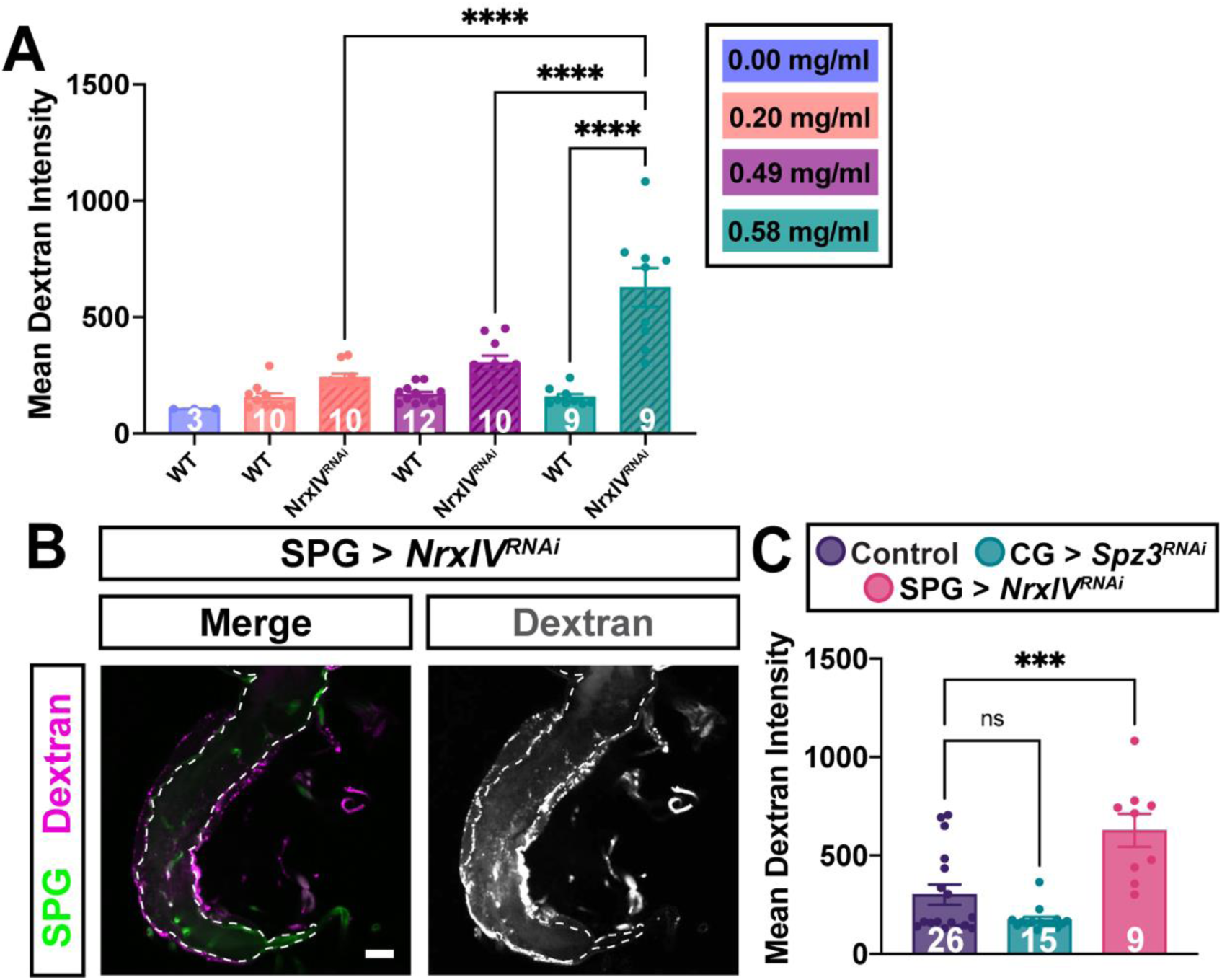
Intensity of Dextran infiltration across the BBB when compromised by NrxIV reduction, related to. Figure 7. (**A**) Quantification of mean Dextran intensity in control (solid bars) and NrxIV RNAi expressed in SPG (stripped bars) L3 CNS subjected to various concentrations of 70kDa Dextran. (**B**) As a positive control, SPG (green) expressing NrxIV^RNAi^ to disrupt the BBB, allow the Dextran (magenta) to infiltrate the L3 CNS. (**C**) Quantification of the mean intensity within the L3 CNS. N = numbers of animals, denoted in each corresponding bar. (n.s.) Not significant. (***) *P* < 0.001. One-Way ANOVA with multiple comparisons.

**Table S1:**
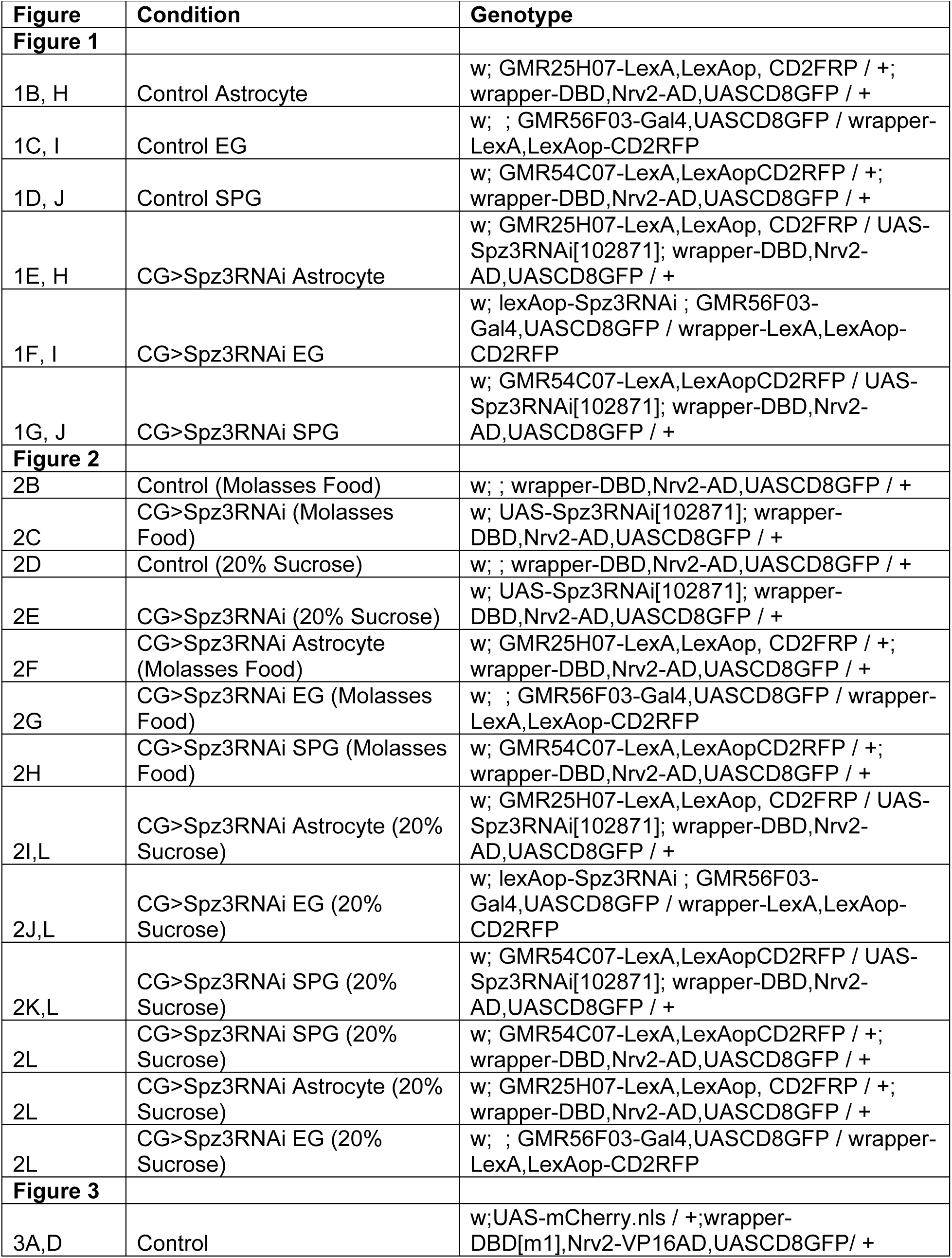
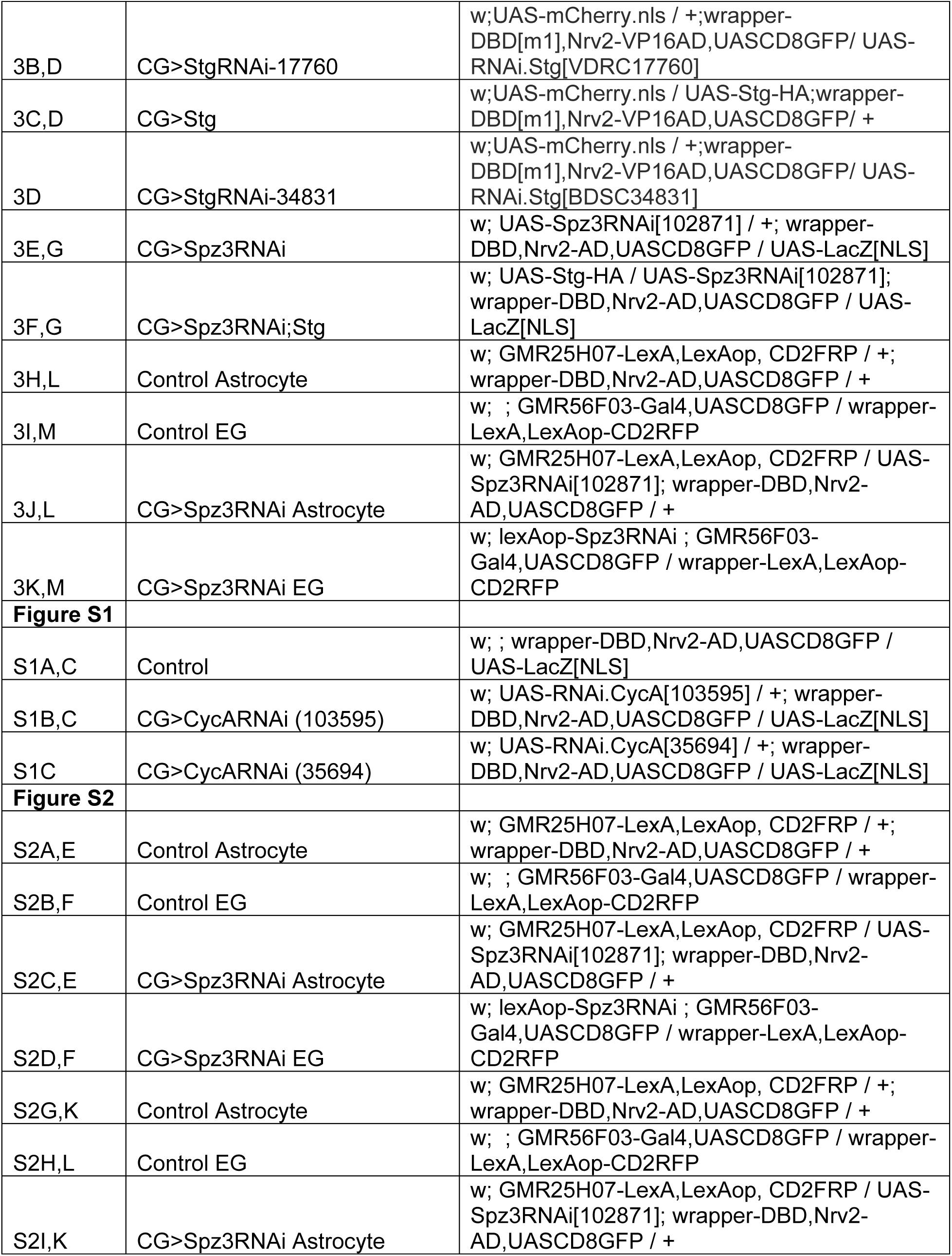
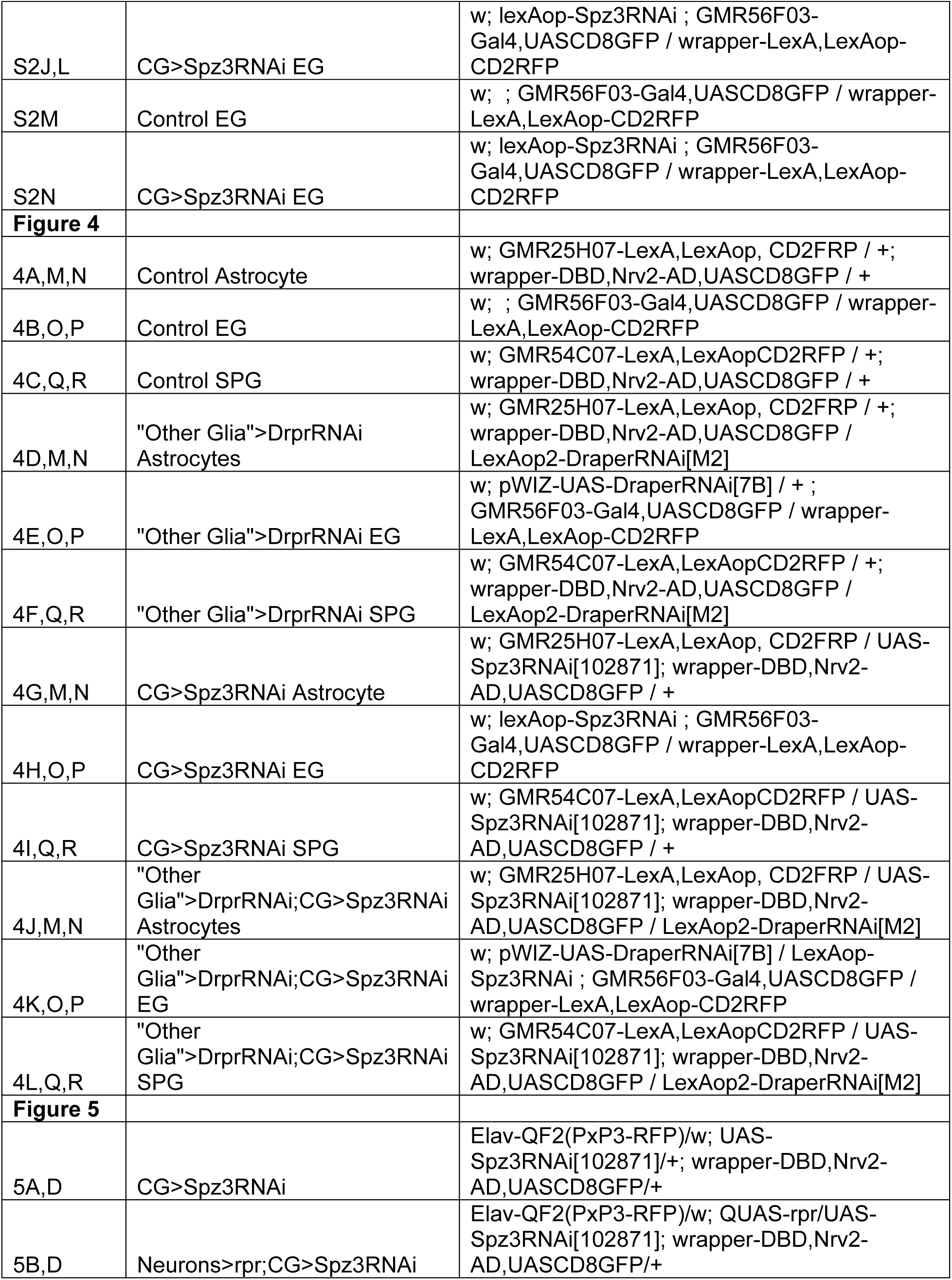
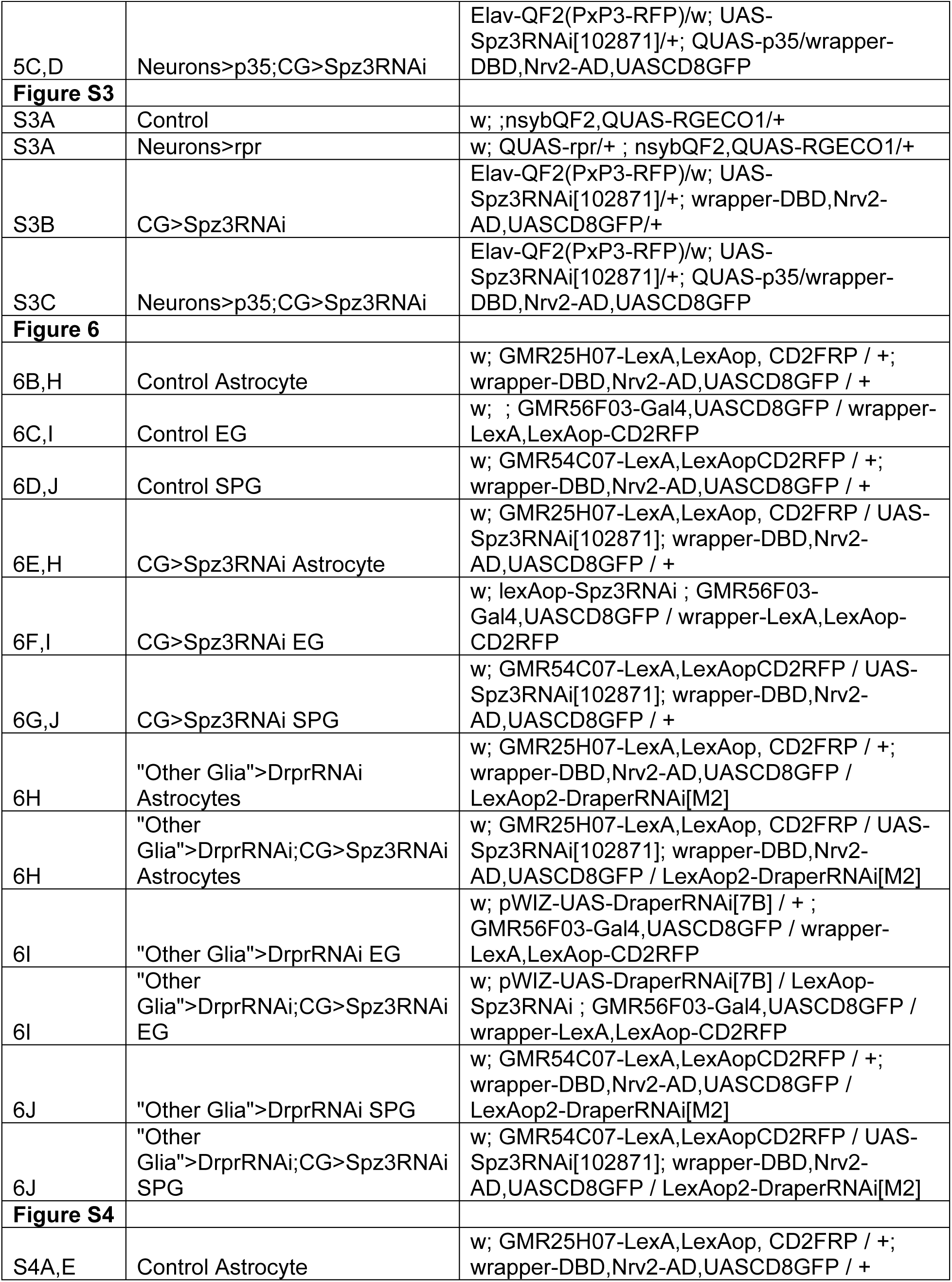
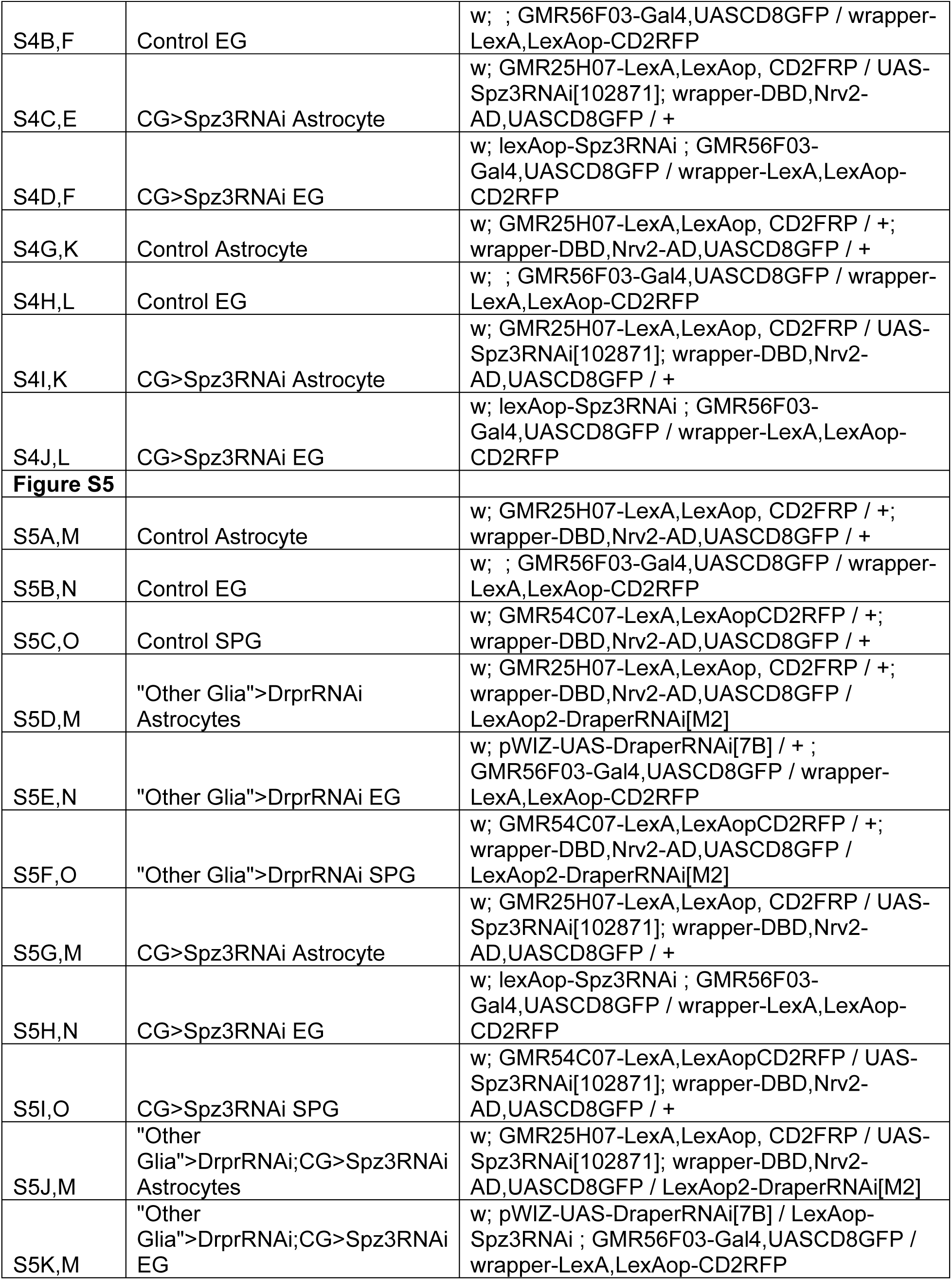
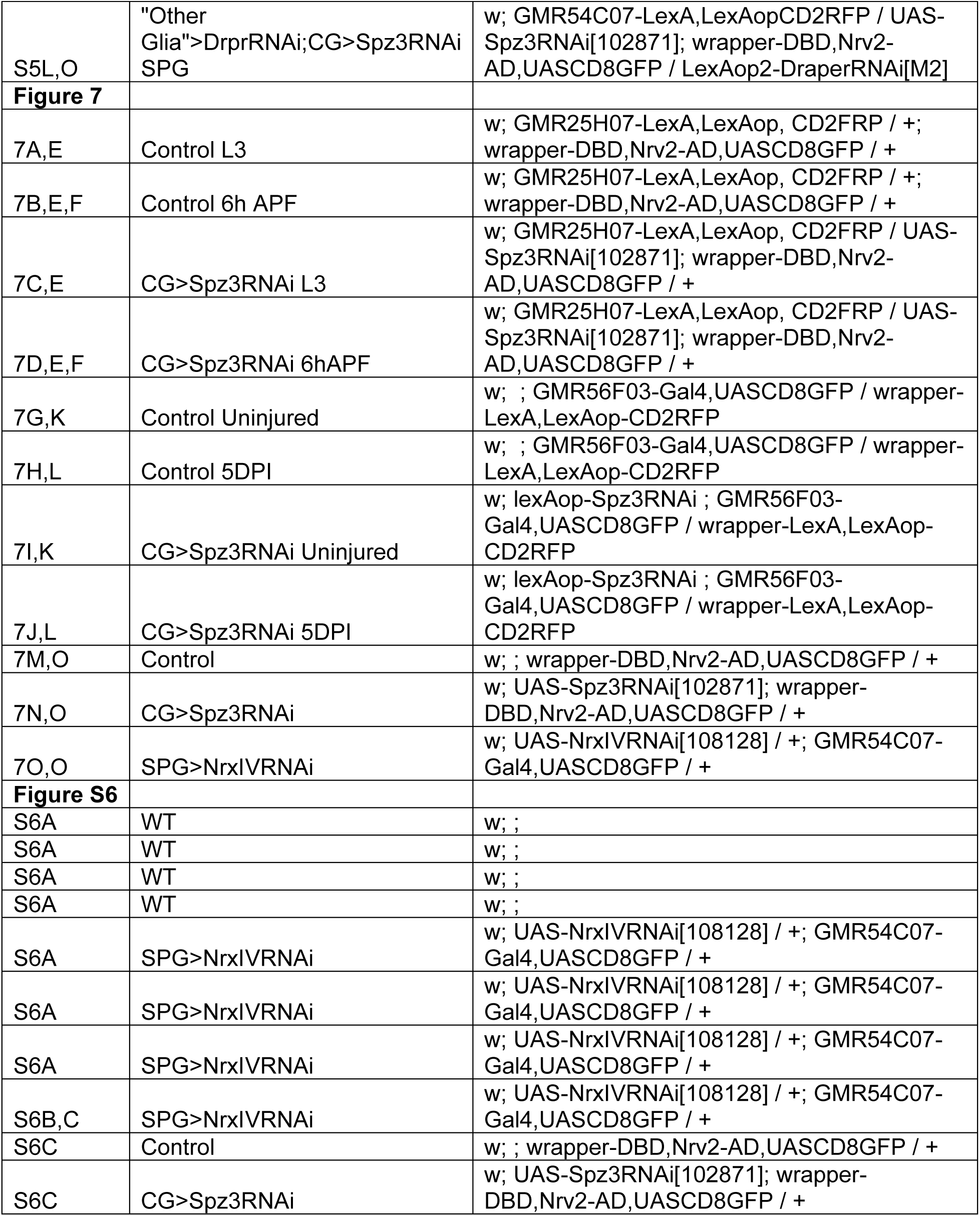
Genotypes for all figures.

## Key resources table

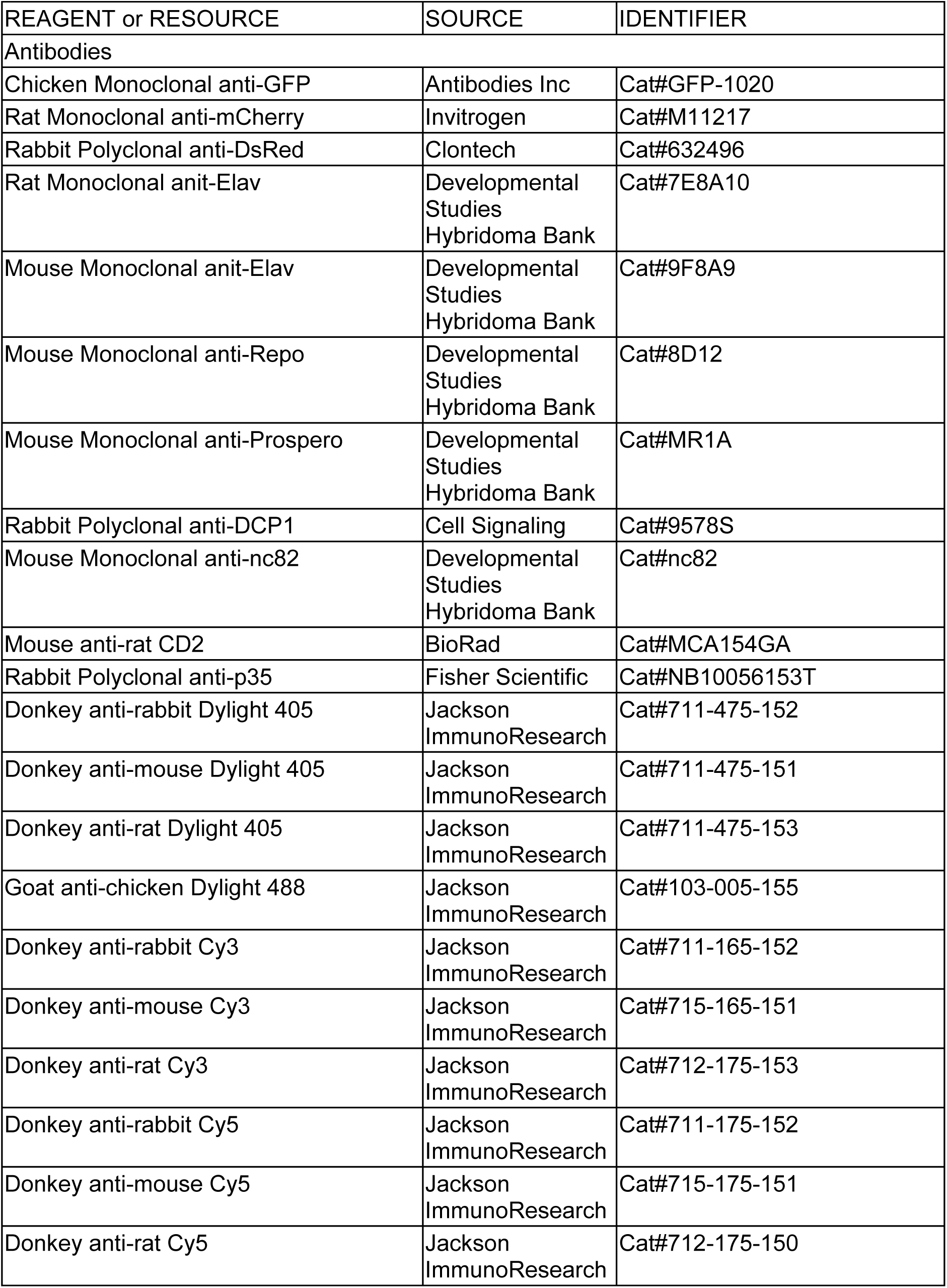
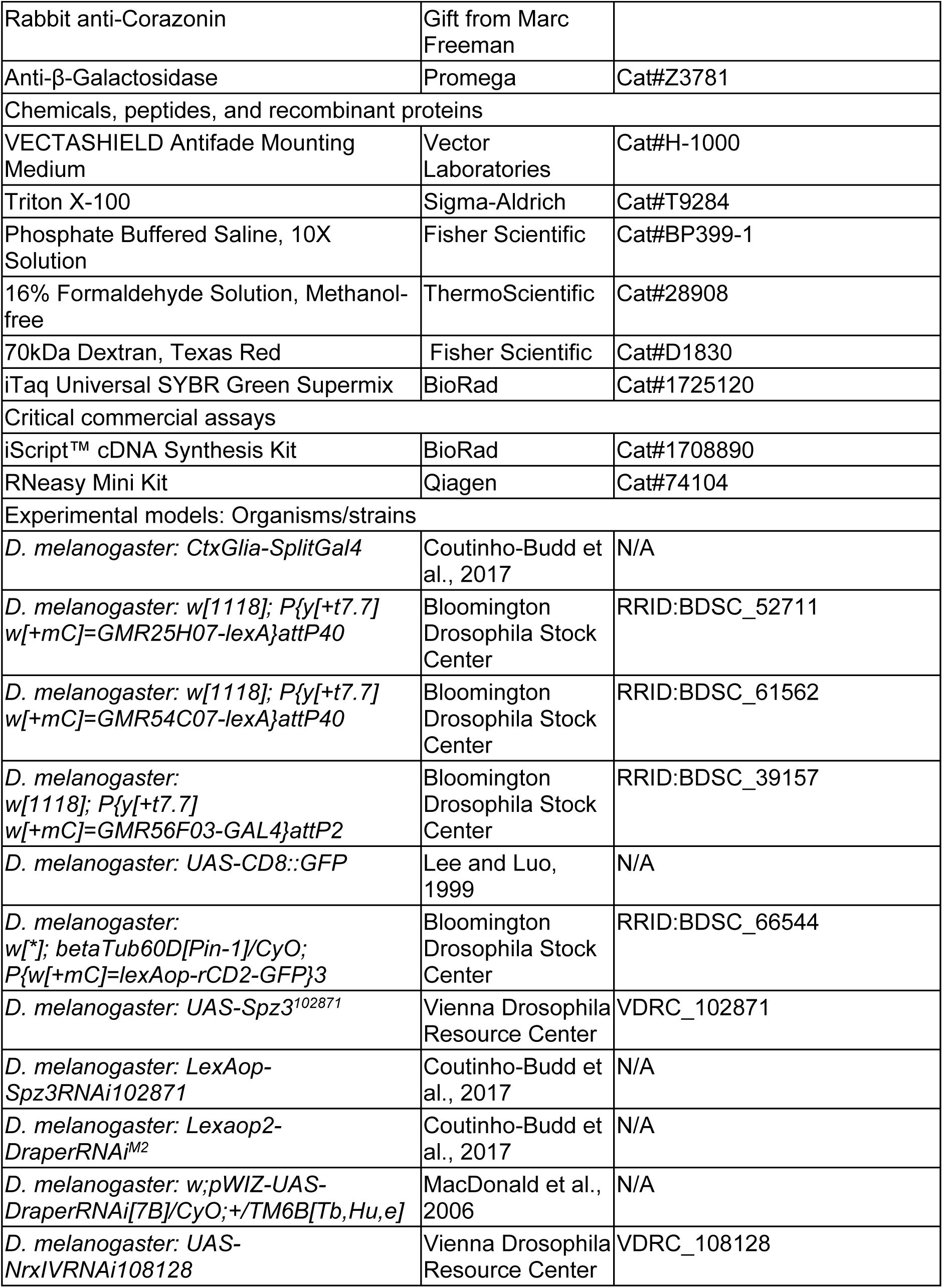
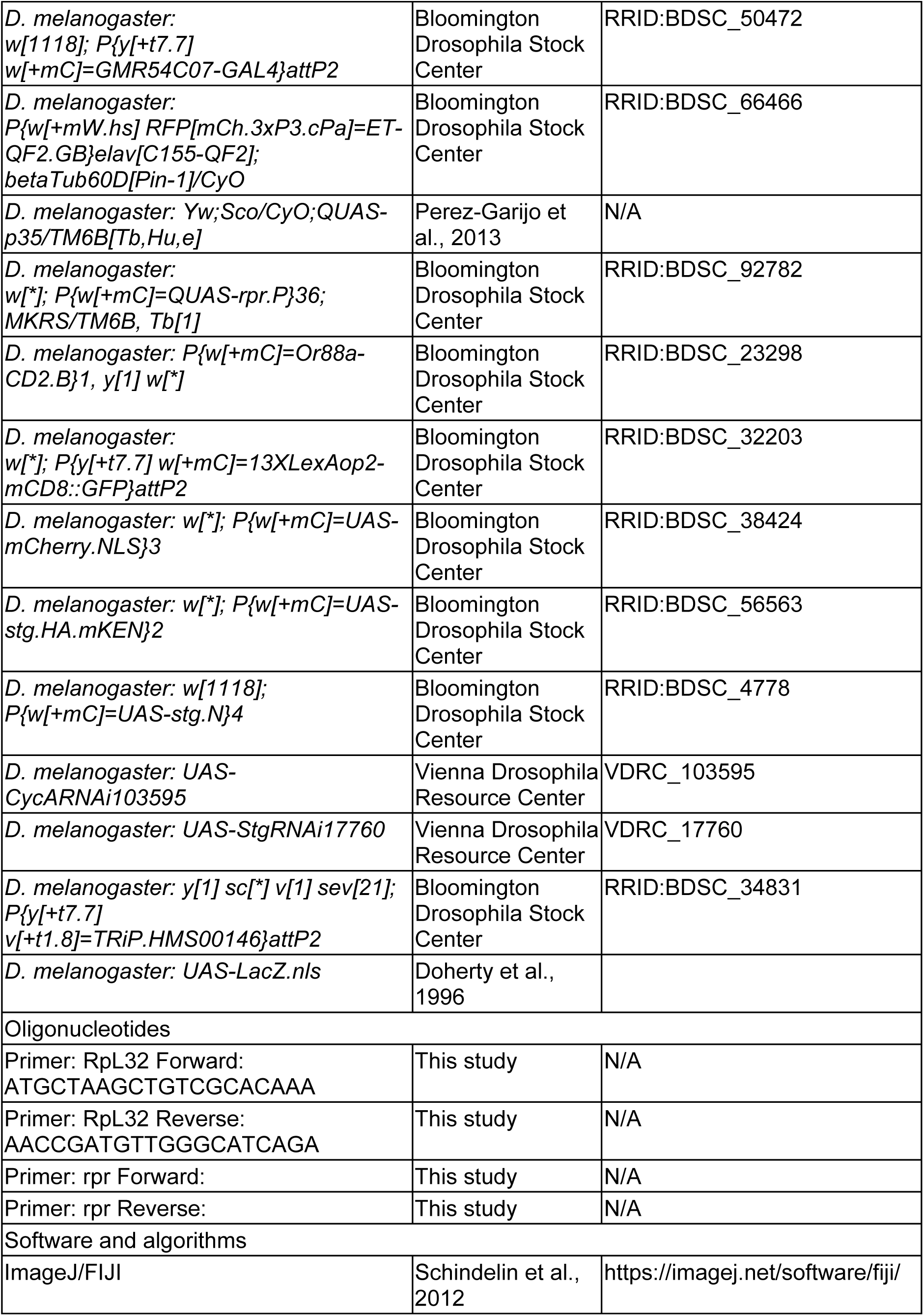
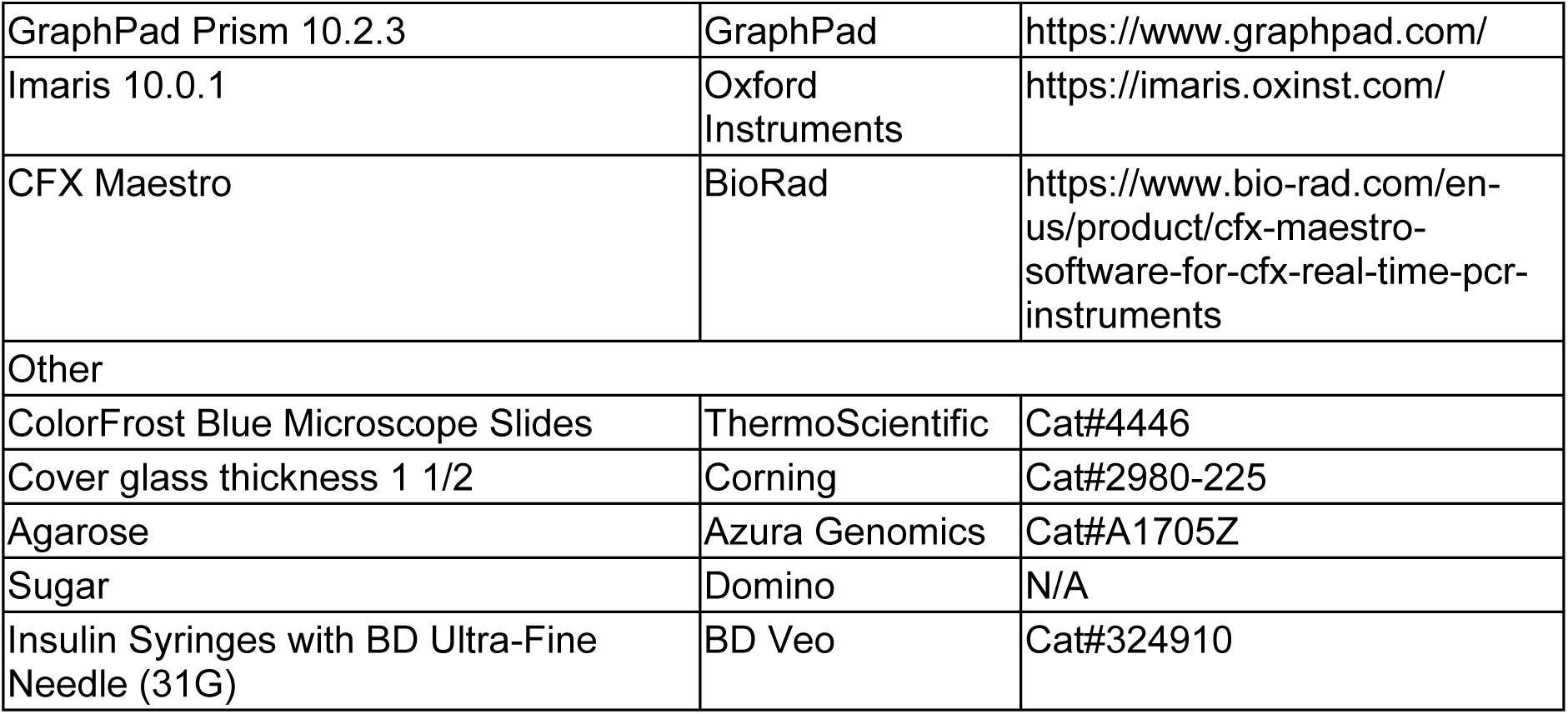

